# Structural Insights into Branch Site Proofreading by Human Spliceosome

**DOI:** 10.1101/2022.11.07.515429

**Authors:** Xiaofeng Zhang, Xiechao Zhan, Tong Bian, Fenghua Yang, Pan Li, Yichen Lu, Zhihan Xing, Qiangfeng Cliff Zhang, Yigong Shi

## Abstract

Selection of the pre-mRNA branch site (BS) by U2 snRNP is crucial to prespliceosome (A complex) assembly. The RNA helicase PRP5 proofreads BS selection; but the underlying mechanism remains unclear. Here we report the atomic structures of two sequential complexes leading to prespliceosome assembly: human 17S U2 snRNP and a cross-exon pre-A complex. PRP5 is anchored on 17S U2 snRNP mainly through occupation of the RNA path of SF3B1 by an acidic loop of PRP5; the helicase domain of PRP5 associates with U2 snRNA; the BS-interacting stem loop (BSL) of U2 snRNA is shielded by the splicing factor TAT-SF1, unable to engage the BS. In the pre-A complex, an initial U2/BS duplex is formed; the translocated helicase domain of PRP5 stays with U2 snRNA; the acidic loop still occupies the RNA path. The pre-A conformation is specifically stabilized by the splicing factors SF1, DNAJC8 and SF3A2. Cancer-derived mutations in SF3B1 damage its association with PRP5, compromising BS proofreading. Together, these findings reveal key insights into prespliceosome assembly and BS selection/proofreading by PRP5.

## Main

Splicing of pre-mRNA is executed by the spliceosome, a dynamic supramolecular machinery ^1,2^. Spliceosome is assembled on pre-mRNA by recognizing conserved elements of an intron: 5’-splice site (5’SS), BS, polypyrimidine tract (PPT), and 3’-splice site (3’SS). First, U1 snRNP, SF1, U2AF2 and U2AF1 recognize 5’SS, BS, PPT and 3’SS, respectively, forming the E complex. Then, U2 snRNP displaces SF1 to allow duplex formation between U2 snRNA and the BS, leading to formation of the A complex ^3–6^. In higher eukaryotes, most genes undergo alternative splicing through selection of different splicing sites ^7–9^. Therefore, formation of the A complex, which selects the BS, is a focal point of splicing regulation ^10,11^. Unfortunately, the steps behind the E-to-A transition are poorly understood.

The RNA helicase PRP5 (also known as DDX46) plays a key role during assembly of the A complex ^4–6,12–14^. In human 17S U2 snRNP, U2 snRNA is unable to pair up with the BS because the relevant sequences of U2 snRNA form the double stranded BSL ^15,16^. ATP hydrolysis by PRP5 leads to the unwinding of BSL, thus promoting formation of the U2/BS duplex. In yeast, Prp5 may proofread the U2/BS duplex because Prp5 mutations allowed usage of cryptic BS and 3’SS ^6,17–19^. However, the mechanism by which PRP5 regulates splicing fidelity remains unclear.

Most known cancer-derived mutations that dysregulate RNA splicing affect residues in the HEAT repeats of SF3B1 ^20–22^. A large majority of these mutations are clustered in the RNA path of SF3B1 ^23,24^. A portion of the corresponding mutations in Hsh155 (yeast orthologue of SF3B1) altered Prp5 interaction and BS selection ^18,19^. Despite these tantalizing clues, how these cancer mutations interfere with normal splicing remains enigmatic.

Addressing these key questions necessitates structural elucidation of intermediate spliceosomal complexes leading to formation of the A complex. In this study, we report the high-resolution cryo-EM structures of human 17S U2 snRNP and a cross-exon pre-A complex arrested by a splicing inhibitor. Together with structure-guided biochemical analysis, our study allows proposition of a working model on proofreading of BS selection and assembly of the A complex.

## Results

### Sample preparation and electron microscopy

Because PRP5 is an integral component of U2 snRNP ^25^, we used a Flag-tagged PRP5 to purify endogenous human 17S U2 snRNP. The purified sample was applied to gradient centrifugation under crosslinking condition ^26,27^. The final sample only contains U2 of the five snRNAs and all known protein components of 17S U2 snRNP (Extended Data Fig. 1a, 1b). 4,273 micrographs were recorded using a K3 detector mounted on a Titan Krios microscope, yielding 1,972,935 particles (Extended Data Figs. 1c and 2). After 2D and 3D classifications, 485,418 particles give a final reconstruction of 17S U2 snRNP at an average resolution of 2.5 Å (Fig. 1a; Extended Data Figs. 2, 3; Extended Data Tables 1, 2). Soft masks on the N-terminal portion of PRP5, TAT-SF1, and the BSL led to improved local resolutions (Extended Data Figs. 2, 3a).

**Fig. 1.**
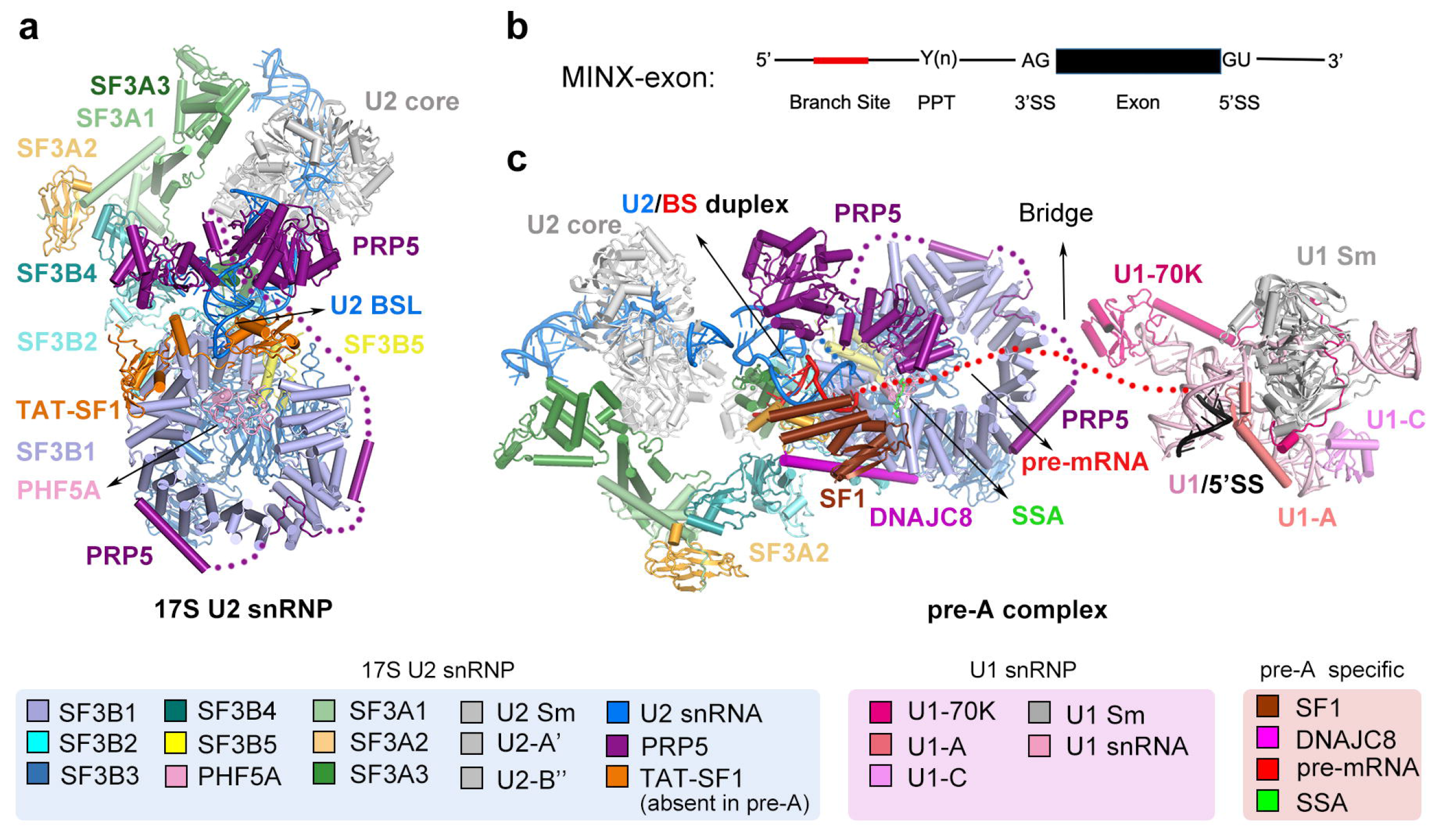
Cryo-EM structures of 17S human U2 snRNP and the human pre-A spliceosome. **a**, Overall structure of 17S human U2 snRNP. The final atomic model includes the SF3b complex, SF3a complex, U2 core (Sm ring, U2-A’, U2-B” and U2 snRNA), the splicing factor TAT-SF1, and the ATPase/helicase PRP5. Protein and RNA components are color-coded and tabulated below the structure. **b**, Schematic diagram of the MINX-exon pre-mRNA. This pre-mRNA allows cross-exon assembly of the human pre-A complex. **c**, Overall structure of the human pre-A complex. The human pre-A complex has a bi-lobal organization, comprising a large U2 snRNP module and a small U1 snRNP module. U2 snRNA forms an initial duplex with the BS. Two pre-A specific factors (SF1 and DNAJC8), but not TAT-SF1, are present in the pre-A complex.

Next, to enrich the intermediate spliceosomal complexes, we used the inhibitor spliceostatin A (SSA), which stalls splicing prior to formation of the A complex ^28^. We used an exon-definition pre-mRNA (designated as MINX-exon) (Fig. 1b), which allows assembly of cross-exon spliceosomes ^29–32^. The purified sample contains pre-mRNA, U1 and U2 snRNAs, and most protein components of the A complex (Extended Data Fig. 4a, 4b). 22,451 electron micrographs were collected, generating 5,156,919 particles (Extended Data Figs. 4c, 4d and 5). Initial data processing revealed presence of a bi-lobal shaped spliceosomal complex (the pre-A complex). Soft masks were separately applied to the two lobes: U2 and U1 snRNPs (Extended Data Fig. 5). The final reconstruction of the human pre-A complex displays an average resolution of 3.0 Å in the U2 snRNP region (Extended Data Figs. 5, 6; Extended Data Tables 1, 3).

The EM maps for 17S U2 snRNP and pre-A complex reveal a number of previously unrecognized features that are functionally important (Extended Data Figs. 7-9). Using a combination of *de novo* modeling and structure docking, we have generated atomic coordinates for these two human complexes (Fig. 1a, 1c; Extended Data Tables 1-3).

### PRP5 is anchored on the RNA path of SF3B1

The 2.5-Å EM map reveals, and allows atomic modeling of, the interface between PRP5 and SF3B1 (Fig. 2a), which has remained elusive despite several published structures of human 17S U2 snRNP ^15,31,33^. Three structural motifs at the N-terminal half of PRP5 closely interact with SF3B1, tethering PRP5 to U2 snRNP. These motifs include an elongated helix α1 (residues 152-178), an acidic loop (residues 195-209), and helix α2 (residues 225-243) that harbors the DPLD-motif ^18,34^ (Fig. 2a).

**Fig. 2.**
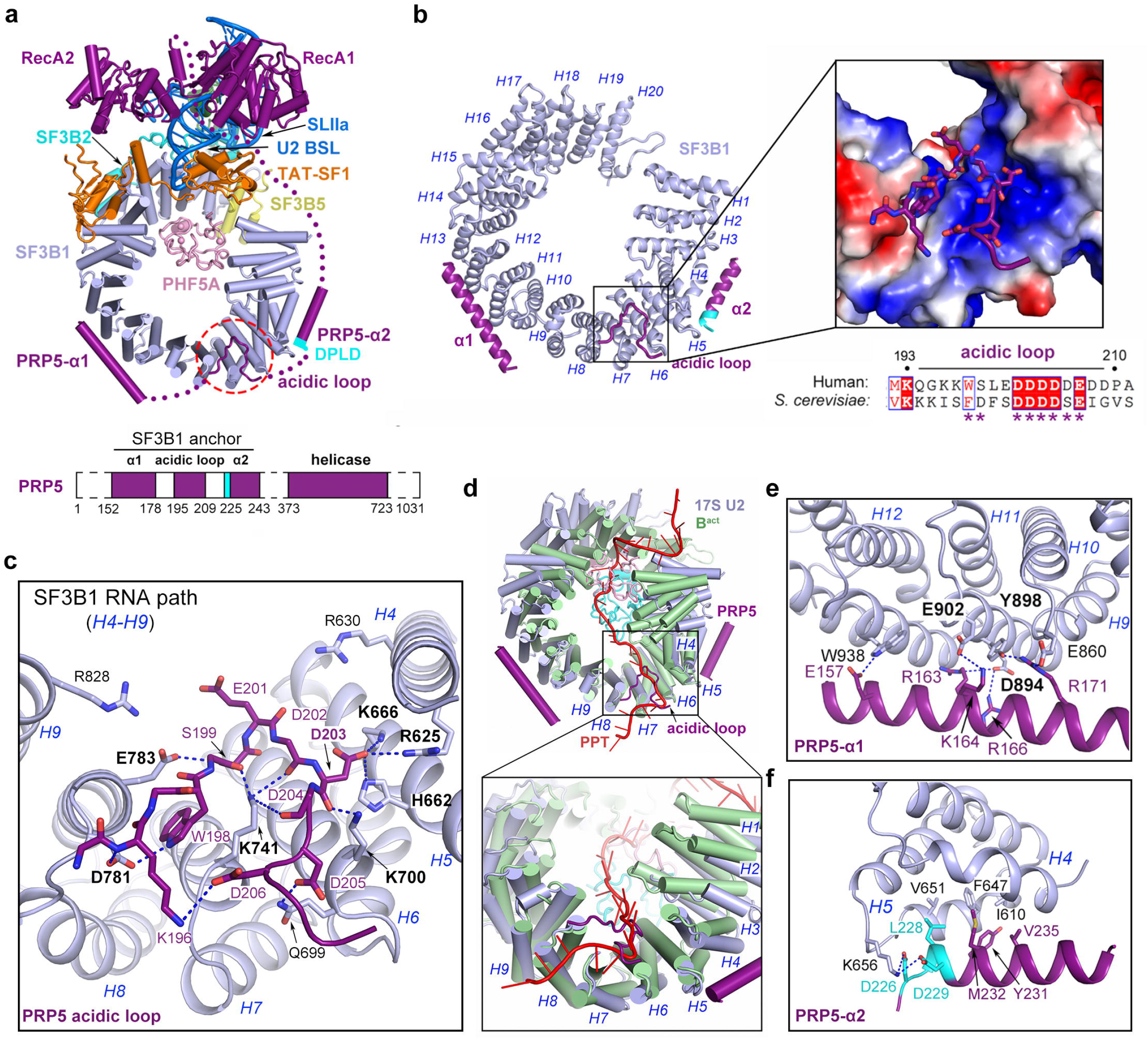
PRP5 is anchored to 17S U2 snRNP through its N-terminal acidic loop. **a**, Overall structure of the core region of 17S U2 snRNP. Only select proteins around the BSL are shown. BSL is sandwiched by TAT-SF1 and the helicase domain of PRP5. PRP5 is anchored on SF3B1 through three motifs in its N-terminal sequences: helix α1, an acidic loop, and helix α2. **b**, An overall view of the interface between the N-terminal sequence of PRP5 and SF3B1. Each HEAT repeat of SF3B1 is indicated by italic H followed by its repeat number. The positively charged RNA path on SF3B1 is occupied by the acidic loop of PRP5 (inset). Sequences of the acidic loop are conserved between *S. cerevisiae* and human (below the inset). Residues that directly interact with SF3B1 are indicated by asterisks. **c**, A close-up view on the interface between the acidic loop of PRP5 and HEAT repeats 4-9 of SF3B1. H-bonds are represented by blue dashed lines. **d**, The positively charged surface of the RNA path is occupied by the PPT of pre-mRNA in the assembled spliceosomes. Shown here is the overlay of SF3B1 from 17S U2 snRNP (light blue) and the human B^act^ complex (green) (PDB code: 5Z56) ^35^. Structural alignment was made using HEAT repeats 4 through 9 (RNA path) of SF3B1. **e**, A close-up view on the interface between PRP5-α1 and HEAT repeats 10-12 of SF3B1. **f**, A close-up view on the interface between PRP5-α2 and HEAT repeats 3-5 of SF3B1. The DPLD motif (cyan) ^5^ interacts with Lys656 from SF3B1.

The most notable feature of the PRP5-SF3B1 interface is placement of the PRP5 acidic loop in the RNA path of SF3B1 (Fig. 2b, 2c), where the polypyrimidine tract (PPT) of pre-mRNA binds in the assembled spliceosomes ^24,35–37^ (Fig. 2d). The RNA path of SF3B1 is heavily enriched by positively charged residues from HEAT repeat 4 (HR4) through HR9, which mediate hydrogen bonds (H-bonds) to negatively charged residues from PRP5 (Fig. 2b). At the center of the interface, Lys741 donates three H-bonds to the carbonyl oxygen atoms of residues 199/202/204 from PRP5 (Fig. 2c). Asp203 from PRP5 accepts three H-bonds from His662, Arg625 and Lys666. At the periphery of the interface, Gln699/Lys700 from HR7 and Asp781/Glu783 from HR8 make H-bonds to Asp205/Asp203 and Trp198/Leu200 from PRP5, respectively (Fig. 2c). Obviously, the acidic loop of PRP5 in the RNA path must be replaced by the PPT of pre-mRNA in the assembled spliceosomes (Fig. 2d). In this sense, PRP5 may safeguard the RNA path from unwanted binding of non-specific RNA before spliceosome assembly.

In addition, helices α1 and α2 of PRP5 also contribute to its association with SF3B1. PRP5-α1 binds the lateral exterior of HR10–HR12 of SF3B1 (Fig. 2b, 2e). This interface features six specific H-bonds. Four positively charged residues Arg163, Lys164, Arg166 and Arg171 of PRP5 donate H-bonds to Asp894, Glu902, Asp894 and Glu860/Tyr898 of SF3B1, respectively (Fig. 2e). Glu157 of PRP5-α1 accepts a H-bond from Trp938 of SF3B1. PRP5-α2 associates with HR4 and HR5 (Fig. 2b, 2f). Asp226 and Asp229 of the DPLD motif from PRP5-α2 accept two H-bonds from Lys656 of HR5. Four residues Leu228/Tyr231/Met232/Val235 of PRP5-α2 make van der Waals contacts to Ile610, Phe647 and Val651 of SF3B1. These structural features are consistent with published biochemical observations ^18^.

### BSL recognition in 17S U2 snRNP

The 5’ sequence of U2 snRNA in human 17S U2 snRNP comprises three elements: stem-loop I (SLI, nucleotides G12 through C21), BSL (U25 through C45), and stemloop IIa (SLIIa, U46 through U65) ^15,16^ (Fig. 3a). The three-way junction of U2 snRNA is stabilized by SF3A3, SF3B2, SF3B1, TAT-SF1, and PRP5. In particular, the Separator helix of SF3A3 is inserted into the space between SLI and SLIIa (Fig. 3a), whereas RecA1 of PRP5 directly contacts SLIIa (Fig. 2a). Our EM reconstruction reveals previously unrecognized structural features of TAT-SF1 that are key to formation of the three-way junction of U2 snRNA (Extended Data Fig. 7b, 7c). The structurally resolved portion of TAT-SF1 comprises an RRM domain (residues 127-220), a UHM domain (residues 264-347), and the intervening Linker domain (residues 235-264) (Fig. 3a).

**Fig. 3.**
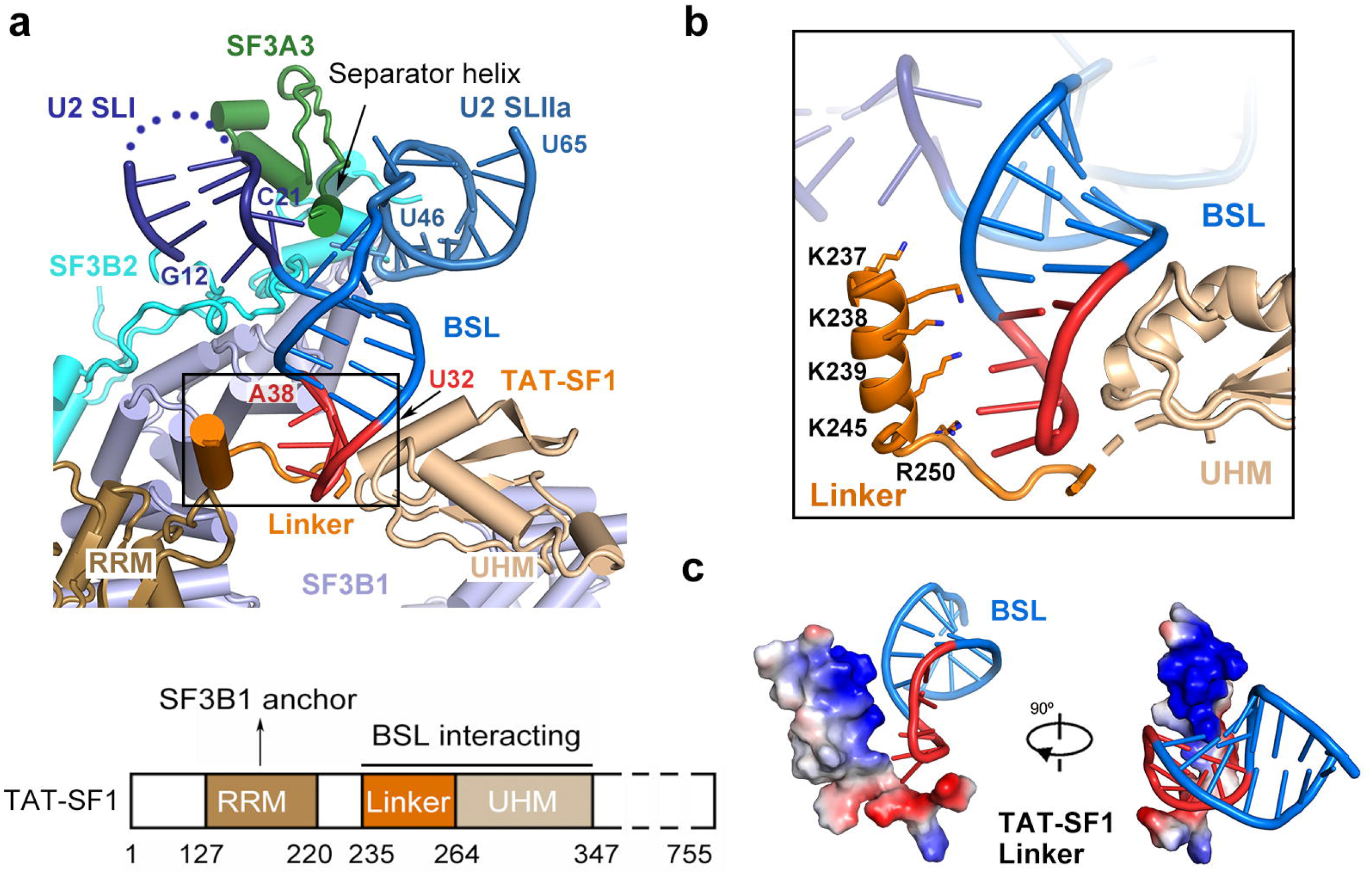
Recognition of the BSL of U2 snRNA. **a**, The three-way junction of U2 snRNA is stabilized by SF3A3, TAT-SF1, SF3B2 and SF3B1. A short α-helix from SF3A3 (known as separator helix ^15^) is inserted into the cleft between SLI and SLIIa and interacts with both RNA strands of BSL. The BSL nucleotides that later form the U2/BS duplex are colored red. The newly identified Linker domain (orange) of TAT-SF1 caps the tip of BSL. SF3B2 and SF3B1 hold the three-way junction from behind. A schematic diagram of the TAT-SF1 sequence features is shown below the structure. **b**, A close-up view on the interface between BSL and the Linker domain of TAT-SF1. A poly-lysine motif from the Linker domain places its positively charged side chains towards the negatively charged phosphate backbone of BSL (upper panel). **c**, Closeup views on the electrostatic surface potential of the TAT-SF1 poly-lysine helix. Blue, positive charges; red, negative charges.

The Linker domain, but not the RRM domain, interacts with the BSL (Fig. 3b). The Linker domain comprises an α-helix and an ensuing loop. The α-helix and the UHM domain are placed on two opposing sides of the double-stranded BSL, with the connecting loop capping the tip of BSL. Four Lys residues 237/238/239/245 and Arg250 from the α-helix constitute a positively charged surface epitope that directly contacts the nucleotides of BSL (Fig. 3b, 3c). This structural organization shields the BSL, making it inaccessible to pre-mRNA. Thus, displacement of the Linker domain is a prerequisite for formation of the U2/BS duplex.

### An initial U2/BS duplex in the human pre-A complex

The human pre-A complex has a bi-lobal organization, with U2 snRNP flexibly connected to U1 snRNP (Fig. 1c). The 3.0-Å reconstruction for U2 snRNP region allows identification of detailed structural features (Extended Data Fig. 6). Compared to 17S U2 snRNP, the conformation of U2 snRNA along with surrounding proteins in the pre-A complex have undergone dramatic rearrangement due to engagement of the pre-mRNA (Fig. 4a). Similar to other SF3B1-targeting inhibitors ^38,39^, SSA is lodged in the Hinged pocket between SF3B1 and PHF5A.

**Fig. 4.**
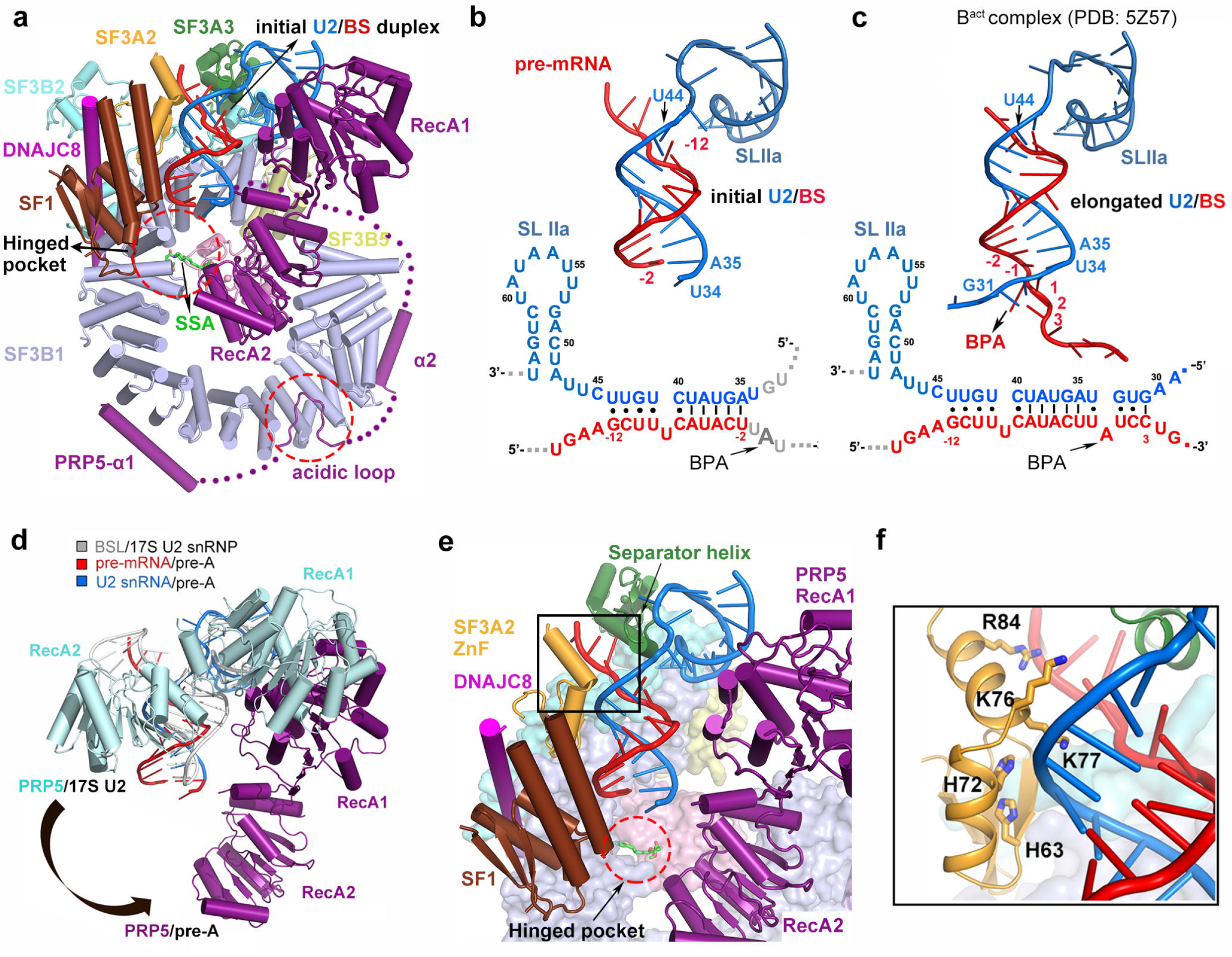
Formation and stabilization of an initial U2/BS duplex in the human pre-A complex. **a**, An overall view of the core of U2 snRNP from the pre-A complex. An initial U2/BS duplex is formed. The splicing inhibitor SSA is lodged in the Hinged pocket between SF3B1 and PHF5A. PRP5 continues to be tethered to SF3B1 through its N-terminal sequences. **b**, Formation of an initial U2/BS duplex in the pre-A complex. A schematic diagram is shown below the carton. Positions relative to the BPA are indicated. The initial U2/BS duplex contains 10 base pairs. **c**, The U2/BS duplex in the B^act^ complex involve 14 base-pairs between pre-mRNA and U2 snRNA. The U2/BS duplex in the A complex should remain the same. **d**, The helicase domain of PRP5 in the pre-A complex (deep purple) is drastically different from that in 17S U2 snRNP (Cyan). **e**, The initial U2/BS duplex is stabilized by surrounding protein components. The U2/BS duplex directly interacts with SF3A3, SF3A2, and SF1. The Separator helix of SF3A3 is inserted into the gap between pre-mRNA and U2 snRNA. SF3A2/SF1 and the helicase domains of PRP5 are placed on two sides of the initial U2/BS duplex. **f**, A close-up view on the interface between the ZnF of SF3A2 and the initial U2/BS duplex.

The most notable feature of the pre-A complex is formation of the U2/BS duplex (Fig. 4a, 4b; Extended Data Fig. 9a). In contrast to 17S U2 snRNP, TAT-SF1 is absent in the pre-A complex. Consequently, the BSL opens up and forms a duplex with pre-mRNA (Fig. 4b). Compared to that in the assembled spliceosomes (Fig. 4c), the length of the U2/BS duplex in the pre-A complex is considerably shorter and hence named the initial U2/BS duplex. Specifically, nucleotides A35 through U44 of U2 snRNA form base-pairs with nucleotides −2 through −12 of pre-mRNA (relative to the position of the branch-point adenosine (BPA)) (Fig. 4b). The BPA and the 5’-end sequences of U2 snRNA are disordered (Fig. 4b; Extended Data Fig. 9a).

Trapping of the pre-A complex is likely a result of SSA binding. In the assembled spliceosomes, the BPA, flipped out of an elongated U2/BS duplex, is specifically recognized by the Hinged pocket between SF3B1 and PHF5A ^24,35,37^. In our pre-A complex, however, SSA occupies the Hinged pocket, making it inaccessible to BPA (Fig. 4a). Consequently, four nucleotides surrounding the BPA form duplex with U2 snRNA only in the assembled spliceosomes (Fig. 4c), but not in the pre-A complex (Fig. 4b). This analysis strongly suggests a coupled relationship between BPA recognition and formation of a stable, elongated U2/BS duplex.

Compared to U2 snRNP, U1 snRNP in the pre-A complex is poorly resolved. With application of a soft mask, the U1 snRNP region displays an average resolution of 15.2 Å (Extended Data Fig. 5), which merely allows docking of the published X-ray structure of U1 snRNP ^40^ into the EM map (Extended Data Fig. 9e). Notably, the PRP5 helix α1 is positioned in close proximity to U1-70K, likely contributing to the connection between U1 and U2 snRNPs (Fig. 1c; Extended Data Fig. 9e). The connection between U1 and U2 snRNPs should also include the single-stranded pre-mRNA, which is presumably flexible (Fig. 1c; Extended Data Fig. 9e).

### PRP5 and stabilization of the initial U2/BS duplex

PRP5 is essential for BSL opening and U2/BS proofreading through unclear mechanisms ^6,12,14,17^. Compared to 17S U2 snRNP, the 3’-sequences of U2 snRNA, including SLIIa and the 3’-strand of BSL, remain nearly unchanged in the pre-A complex (Extended Data Fig. 10a). In contrast, the 5’-sequences of U2 snRNA, including SLI and the 5’-strand of BSL, have become disordered in the pre-A complex. The space vacated by SLI and the 5’-strand of BSL is occupied by the BS, which forms the initial U2/BS duplex with the 3’-strand of BSL (Extended Data Fig. 10a). The RNA changes are accompanied by rearrangement of surrounding proteins, in particular PRP5. Relative to that in 17S U2 snRNP, RecA2 of PRP5 in the pre-A complex undergoes a rotation of about 50 degrees and a translocation of over 50 Å (Fig. 4d). Compared to RecA2, RecA1 displays a considerably smaller positional shift.

Similar to that in 17S U2 snRNP, RecA1 of PRP5 continues to associate with SLIIa of U2 snRNA in the pre-A complex (Fig. 4e). The EM density for the very 5’-end of U2 snRNA can be traced to a location between RecA1 and RecA2 (Extended Data Fig. 10b), suggesting that the 5’-end of U2 snRNA may still be bound to PRP5. These observations suggest a model in which PRP5 pulls the 5’-sequences of U2 snRNA to unwind the BSL, with RecA1 to serve as the pivot for RecA2 to exert the force and translocate on single-stranded U2 snRNA.

In addition, the initial U2/BS duplex is surrounded by SF3A3, the zinc finger (ZnF) of SF3A2, and the splicing factor SF1 (Fig. 4e). Compared to that in 17S U2 snRNP, SF3A3 maintains the same overall conformation, except that the Separator helix is now inserted into the space between pre-mRNA and U2 snRNA. The ZnF of SF3A2, which is disordered in 17S U2 snRNP, closely associates with the initial U2/BS duplex in the pre-A complex. Five positively charged residues Arg84/Lys76/Lys77/His72/His63 from SF3A2-ZnF contact U2 snRNA (Fig. 4f). These observations suggests a crucial role of SF3A2-ZnF in stabilizing the U2/BS duplex during assembly of the A complex, consistent with prior biochemical observations ^41^.

The BS is recognized first by SF1 in the E complex and then by U2 snRNP in the A complex ^42,43^. SF1 was reported to interact directly with protein components of U2 snRNP ^44^, suggesting a transitory role for SF1 in handing over the BS. The pre-A complex presumably captures such an intermediate state, as reflected by the interaction between SF1 and the phosphate backbone of BS (Fig. 4e; Extended Data Fig. 9d). The X-ray structure of SF1 ^42,45,46^ can be readily docked into the EM density map (Extended Data Fig. 9d). Presumably, SF1 facilitates assembly of the A complex by placing BS in the vicinity of U2 snRNP and remains bound until maturation of the A complex ^47^.

### DNAJC8 in the pre-A complex

Formation of the A complex is a multi-step, dynamic process that requires specific splicing factors ^4,48^. Our structure identifies DNAJC8 as a key component of the pre-A complex. Despite the exact role in spliceosome remains unclear, DNAJC8 has been thought to be a molecular chaperone of SRPK1, a SR protein-specific kinase that regulates spliceosome assembly and alternative splicing ^49,50^. The structurally resolved region of DNAJC8 (residues180-233) forms an elongated, curved α-helix (Fig. 5a; Extended Data Fig. 9c). The N-terminal portion of the α-helix interacts with SF1 and SF3A2, likely stabilizing their interaction with U2/BS (Fig. 5a). Glu213 and Arg219 from the middle portion of the helix form H-bonds with Arg1053 and Asp1092 of SF3B1, respectively (Fig. 5b). Trp223 and Phe226 from the C-terminal portion of the α-helix make van der Waals contacts to hydrophobic residues from SF3B1 and SF3B2 (Fig. 5c). Together, these interactions glue DNAJC8 to the periphery of U2 snRNP in the pre-A complex. Notably, DNAJC8 binds to SF3B1 on the opposite site of the Hinged pocket (Fig. 5d). Docking of BPA into the Hinged pocket triggers dramatic conformational change in SF3B1, which presumably disrupts DNAJC8 binding. Release of DNAJC8 in turn may destabilize SF1 binding.

**Fig. 5.**
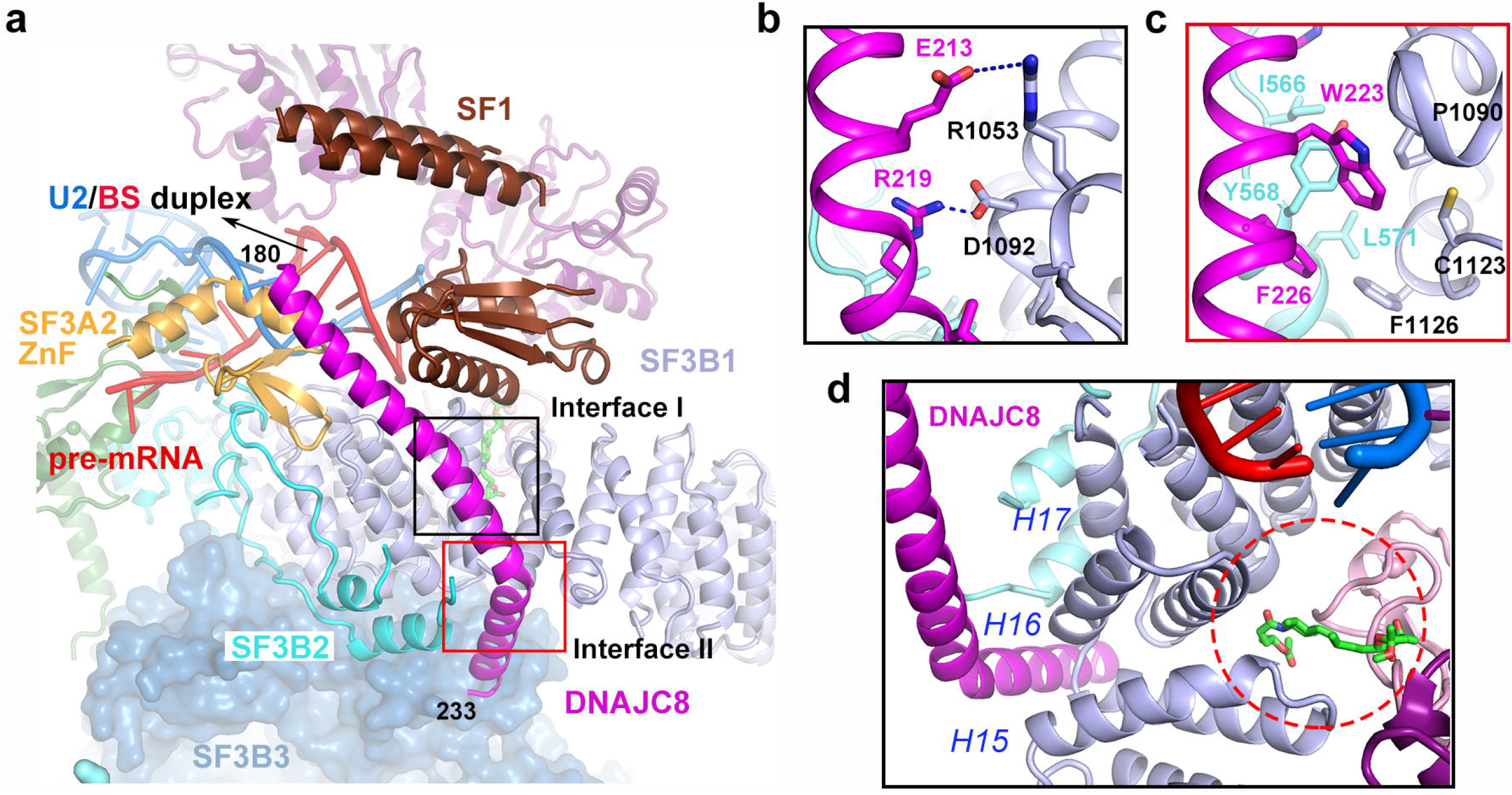
DNAJC8 in the pre-A complex. **a**, The structurally resolved region of DNAJC8 (residues 180-233) forms an elongated and curved α-helix in U2 snRNP. The N-terminal portion of the helix interacts with SF1 and SF3A2; the C-terminal portion binds to SF3B1 and SF3B2. **b**, A close-up view on the interface between the C-terminal portion of the DNAJC8 helix and SF3B1. H-bonds are indicated by blue dashed lines. **c**, A close-up view on the interface between Trp223/Phe226 of the DNAJC8 helix and a hydrophobic surface formed by residues from SF3B1 and SF3B2. **d**, A close-up view on the binding of the inhibitor SSA in the Hinged pocket. The Hinged pocket is formed by HEAT repeats 15-17 of SF3B1. DNAJC8 and SSA are located on two opposing sides of the Hinged pocket.

### Cancer mutations in SF3B1 cripple PRP5 interaction

Cancer-derived mutations have been found in a number of spliceosomal components, in particular SF3B1 ^10,20,51^. These mutations are clustered in the RNA path and surrounding areas, which coincide with the PRP5-binding site on SF3B1 (Fig. 6a). In the RNA path, Asp781 is recurrently mutated in myelodysplastic syndrome (MDS)^52^. Arg625, His662, Lys666, and Lys700, each of which directly interacts with the acidic loop through H-bonds (Fig. 2c), are mutated with high frequency in both MDS and chronic lymphocytic leukemia (CLL) ^20^. Outside the RNA path, Asp894 and Glu902 interact with PRP5-α1; the mutations D894H and E902K are frequently found in patients with CLL ^23,52^.

**Fig. 6.**
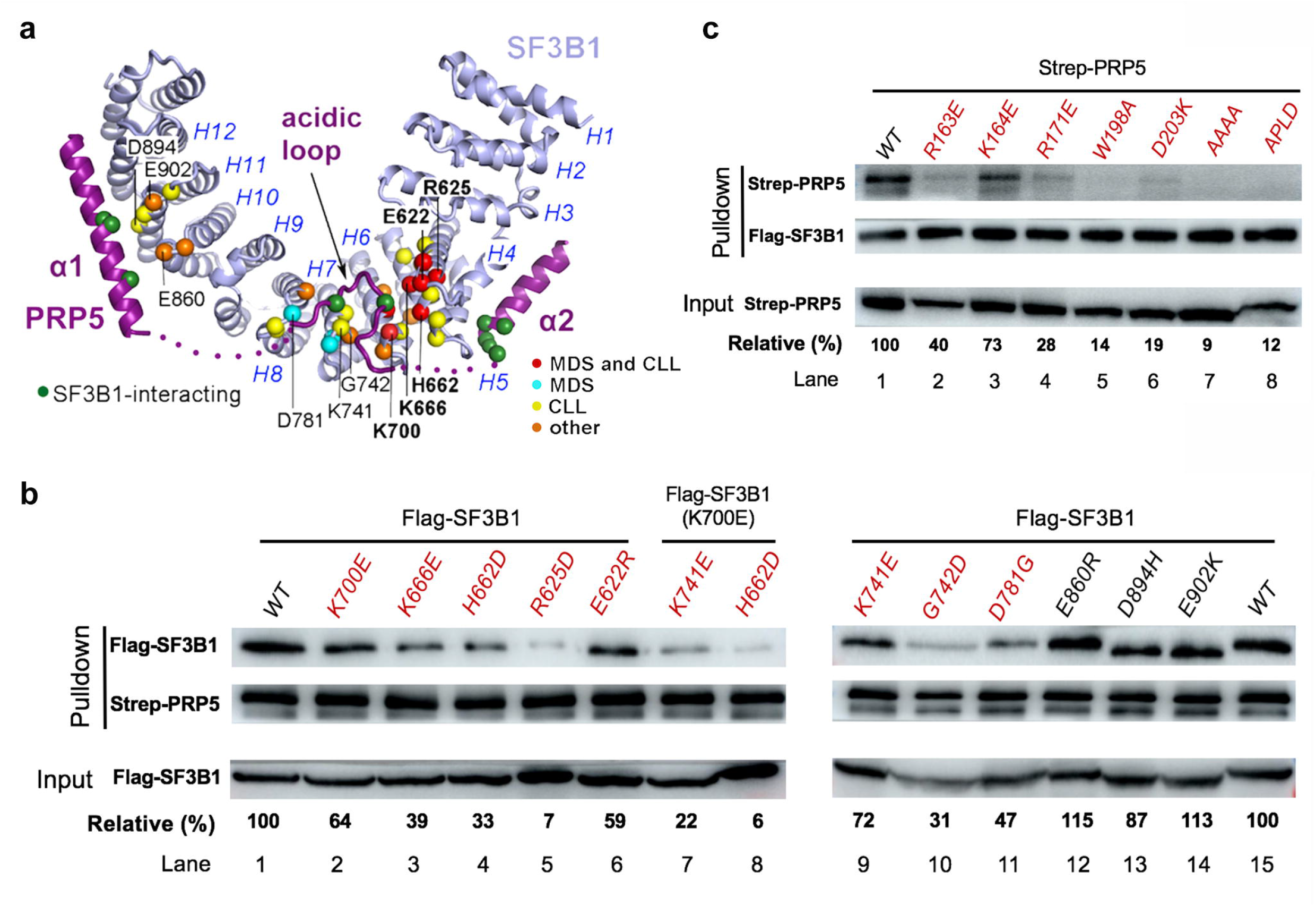
Cancer-derived mutations in the RNA path of SF3B1 impair its interaction with PRP5. **a**, The vast majority of cancer-derived mutations in SF3B1 are clustered at its interface with PRP5. Color coding for mutations: red, both myelodysplastic syndrome (MDS) and chronic lymphocytic leukemia (CLL); cyan, MDS only; yellow, CLL only; orange, other cancers. The five most frequently occurring SF3B1 mutations are highlighted in bold. PRP5 residues that interact with SF3B1 are identified by green spheres. **b**, Cancer-derived mutations in the RNA path of SF3B1 impair its interaction with PRP5. Recombinant Strep-PRP5 was individually incubated with several cell lysates, each containing a distinct SF3B1 mutant. Proteins were selected by Strep-Tactin resin and analyzed by Western blots. The pulldown efficiency, expressed as a ratio of pulldown over input, was averaged for three independent repeats and normalized to the corresponding WT protein. **c**, Mutations in PRP5 cripple its binding to SF3B1. The purified recombinant protein Strep-SF3B1 was incubated individually with cell lysates of different PRP5 mutants and analyzed similarly as above.

To examine the consequences of these mutations, we performed structure-guided biochemical assay. First, wild-type (WT) Strep-tagged PRP5 was used to pull down Flag-tagged SF3B1 (Fig. 6b). As anticipated, WT PRP5 stably interacted with WT SF3B1; the ratio of WT SF3B1 that formed a complex with WT PRP5 over the input was normalized as 100 percent (Fig. 6b, lane 1). Compared to WT SF3B1, the mutation K700E results in a reduced relative binding ratio of 64 percent (Fig. 6b, lane 2). Similar to K700E, all seven other cancer-derived mutations (E622R, R625D, H662D, K666E, K741E, G742D, and D781G) in the RNA path negatively affected its interaction with WT PRP5 (Fig. 6b, lanes 3-6 & 9-11). The double mutant K700E/K741E or K700E/H662D in SF3B1 crippled its interaction with WT PRP5 (Fig. 6b, lanes 7 & 8).

Compared to mutations in the RNA path, mutations in HR10-HR12 of SF3B1, which affect the interface with PRP5-α1, had no impact on the SF3B1-PRP5 interaction (Fig. 6b, lanes 12-14). This result supports a dominant role by the acidic loop of PRP5. To further corroborate this conclusion, we generated seven PRP5 mutants, each carrying one or more missense mutations at the interface with SF3B1 (Fig. 6a). In all cases, these PRP5 mutants exhibited severely diminished binding to WT SF3B1 (Fig. 6c, lanes 2-8). The generally more severe phenotypes by these PRP5 mutations are likely caused by the fact that each PRP5 residue usually interacts with two or more SF3B1 residues. For example, the mutation D203K in PRP5 may abrogate H-bonds with Arg625, His662, and Lys666 of SF3B1; but the mutation H662D in SF3B1 only affects one H-bond with Asp203 (Fig. 2c).

Splicing abnormality has been observed in various types of SF3B1-mutated tumors ^53,54^. Aberrant usage of 3’SS has been established as the most frequent splicing defect and is attributed to altered BS selection ^19,53–56^. Our structural and biochemical observations support the hypothesis that these SF3B1 mutations may compromise BS proofreading due to weakened binding to PRP5 ^18^. To assess this hypothesis, we examined whether reduced PRP5 expression would phenocopy the splicing defects of SF3B1 mutations. Using siRNA knockdown (KD) approach (Extended Data Table 4), we reduced the expression level of PRP5 in HEK293 cells (Extended Data Fig. 11a) and extracted the RNA for transcriptome analysis. Differential analysis of splicing junctions using JuncBASE ^57^ reveals marked differences between the control and *PRP5-KD* cells (Extended Data Fig. 11b, 11c). Among 2,397 aberrantly spliced junctions identified, 1,048 (44%) were novel variants as they had no ensemble transcript identifiers (Extended Data Table 5). A large proportion of aberrant splicing events in *PRP5*-KD cells are intron retention (38%), more than alternative 3’SS (22%) (Extended Data Fig. 11d), which is different from that observed in tumors harboring SF3B1 mutations ^58^. This generally more severe phenotypes in *PRP5*-KD cells could be partially explained by the fact that SF3B1 mutations are heterozygous in tumor cells ^20^. Thus, *PRP5*-KD may undermine the ability of U2 snRNP to engage the BS, leading to intron retention. In addition, reduced expression of PRP5 may compromise BS proofreading, resulting in altered BS and 3’SS selection. Notably, many aberrantly spliced gene isoforms that are frequently found in hematopoietic cancers appear to be sensitive to PRP5 knockdown (Extended Data Fig. 11e; Extended Data Table 5).

These genes exemplified by *ANKHD1* undergo alternative splicing using cryptic 3’SS (Extended Data Fig. 11f). Taken together, our RNA-seq analysis reveals both similarities and differences on altered splicing pattern between SF3B1-mutated tumors and *PRP5*-KD cells, suggesting sophisticated roles of splicing dysregulation in carcinogenesis.

### A working model of BS recognition and proofreading

Structural analysis of human 17S U2 snRNP and the pre-A complex suggests a working model for BS recognition and proofreading. Although the RNA helicase domain of PRP5 is positioned quite differently in these two complexes, the acidic loop and two surrounding helices are anchored on SF3B1 identically (Fig. 7a, 7b). Notably, PRP5-bound SF3B1 exists in an open conformation (Extended Data Fig. 12a). In contrast, SF3B1 adopts a closed conformation in the assembled spliceosomes (Extended Data Fig. 12b-d), in which the PPT, but not the acidic loop of PRP5, is accommodated in the RNA path ^24,35,37^ (Figs. 2d and 7c).

**Fig. 7.**
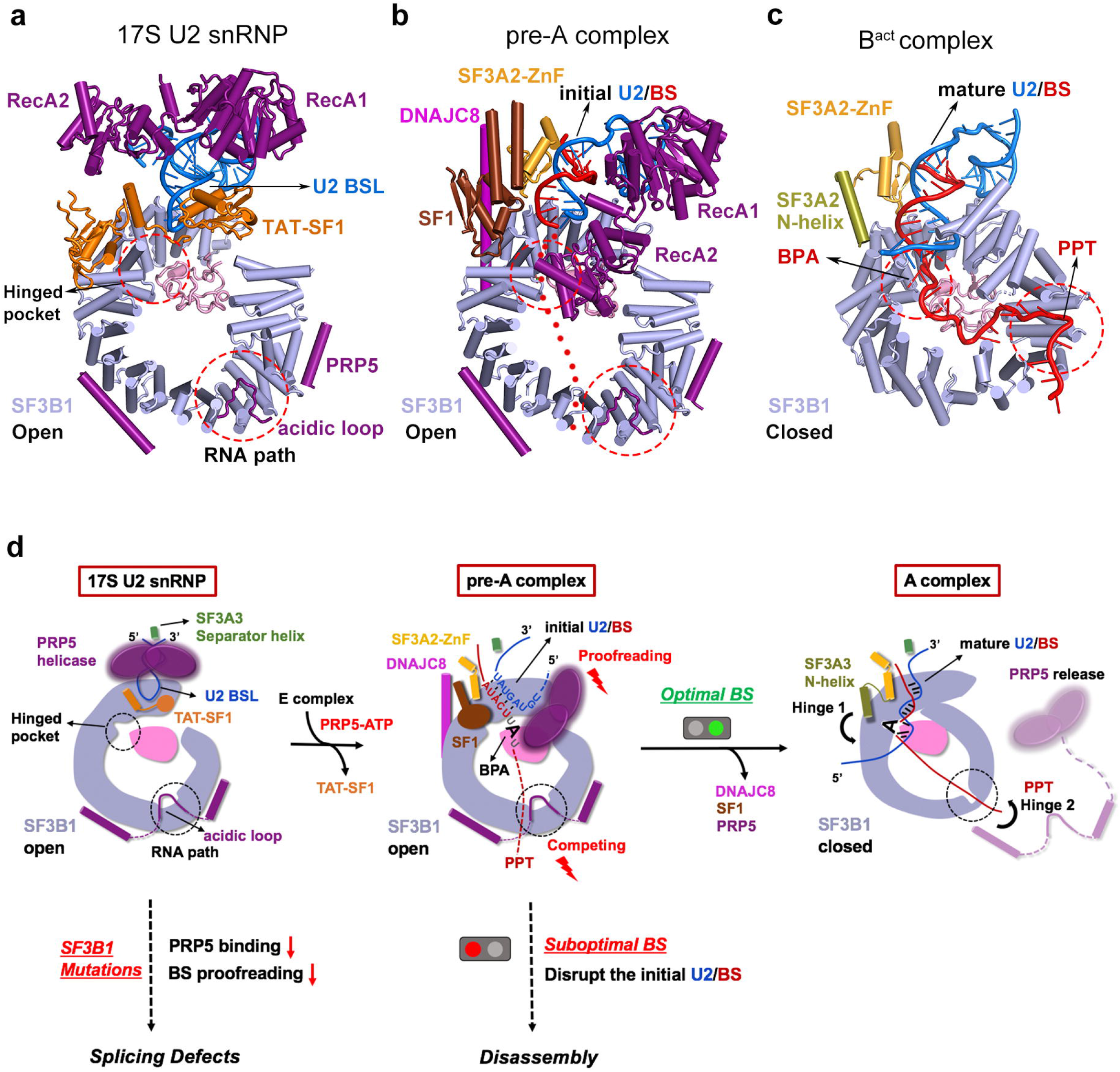
A working model for assembly of the A complex and PRP5-mediated proofreading of BS. **a**, The three-way junction of U2 snRNA is maintained by selected components of human 17S U2 snRNP. The Hinged pocket and the RNA path of SF3B1 are indicated by red dashed circles. **b**, An initial U2/BS complex is stabilized by selected components of the human pre-A complex. Sequences downstream of the BS, including PPT-3’SS and the exon, are indicated by dashed lines. The acidic loop of PRP5 occupies the RNA path, making it inaccessible to pre-mRNA. **c**, A mature U2/BS complex is accommodated by selected components of U2 snRNP core in the human B^act^ spliceosome ^35^. The conformation of U2 snRNP core should remain unchanged between the A and B^act^ complexes. The BPA is positioned in the Hinged pocket and the PPT occupies the RNA path. SF3B1 adopts a closed conformation. The N-helix of SF3A2 is in the same location as SF1/DNAJC8 in the pre-A complex. **d**, A working model on assembly of the A complex and PRP5-mediated proofreading of BS.

During assembly of the A complex, 17S U2 snRNP is brought into the vicinity of the BS via interaction with SF1 ^42,44^. However, due to protection by TAT-SF1, the BSL is inaccessible to pre-mRNA (Figs. 7a). Powered by ATP binding and hydrolysis, PRP5 likely pulls the 5’-end of U2 snRNA to unwind the double-stranded BSL, disrupting its interactions with TAT-SF1 (Fig. 7d). The opened BSL sequences in turn pair up with the BS to form an initial U2/BS duplex. In this intermediate state, SF1 is yet to be completely dissociated from BS. Instead, it binds the initial U2/BS duplex together with SF3A3, SF3A2-ZnF and the helicase domain of PRP5. DNAJC8 specifically interacts with SF3B1 on the opposite side of the Hinged pocket (Fig. 7d).

Although the mechanism of BS proofreading by PRP5 remains unclear ^6,17,18,47^, our structures of 17S U2 snRNP and pre-A complex suggest a model in which PRP5 proofread the initial U2/BS duplex through a competition mechanism. In the pre-A complex, the helicase domain of PRP5 closely interacts with the initial U2/BS duplex, while the acidic loop of PRP5 occupies the RNA path, making it inaccessible for pre-mRNA. An initial U2/BS duplex involving a suboptimal BS has imperfect base-pairing interactions and is thus more likely to succumb to the activity of PRP5, leading to dissociation of the suboptimal BS from U2 snRNA. Conversely, an initial U2/BS duplex involving an optimal BS is less likely to be destabilized by PRP5, thus giving the downstream PPT and 3’SS sequences an improved opportunity to outcompete the acidic loop of PRP5 from the RNA path, ultimately leading to PRP5 dissociation. Concomitant with PRP5 release, BPA is docked into the Hinged pocket and SF3B1 turns into a closed conformation (Fig. 7d).

## Discussion

While this manuscript was under preparation, a structure of the pre-A complex from *S. cerevisiae* was reported at an average resolution of 5.9 Å ^59^. These two complexes are quite different on a number of important criteria. First, the origin of the pre-A complex is yeast in the published study and human in this manuscript. Second, the nature of the pre-A complex is different: intron-definition for yeast ^59^ and exon-definition for human. Third, the relative orientations of U1 and U2 snRNPs are different, resulting in different interfaces (Fig. 8). Importantly, the cross-exon pre-A complex needs to be converted to cross-intron complex to allow progression of splicing reaction; this process requires dramatic remodeling of the spliceosome (Extended Data Fig. 13). Fourth and significantly, there are a number of functionally important and conserved structural features in the human complex that are absent in the yeast structure. For example, the N-terminal half of PRP5 was unresolved in the yeast pre-A complex. Our structure of human 17S U2 snRNP and pre-A complex reveal a key role for the acidic loop of PRP5 in assembly of the A complex and proofreading of the BS. In addition, the splicing factors SF1 and DNAJC8, which stabilize the U2/BS duplex in human pre-A complex, are absent in the yeast complex (Fig. 8).

**Fig. 8.**
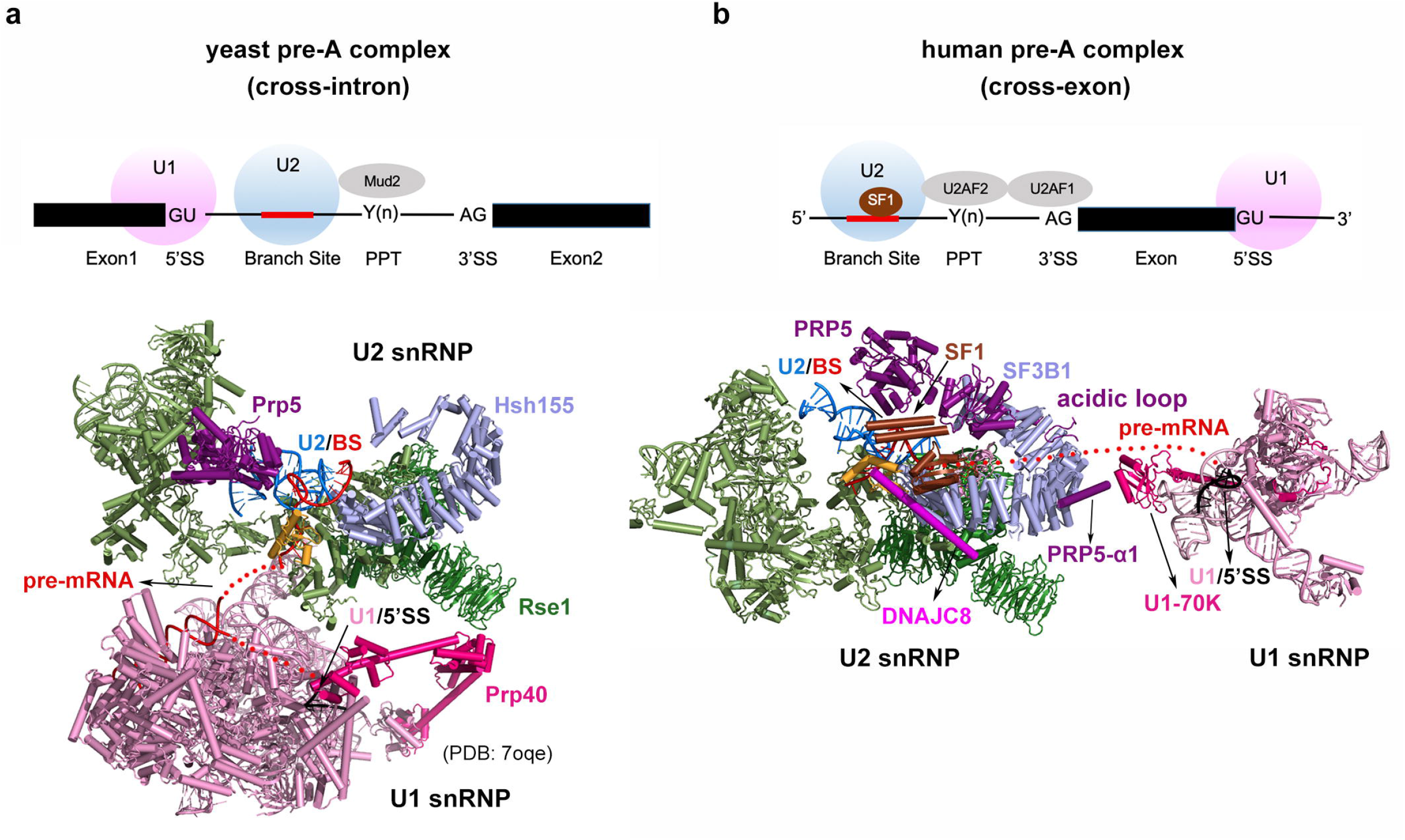
Structural comparison between the yeast intron-defined pre-A complex and the human exon-defined pre-A complex. **a**, Structure of the pre-A complex from *S. cerevisiae* ^59^. The yeast complex was assembled on an intron-definition pre-mRNA. A schematic diagram of the pre-mRNA and its recognition by the pre-A complex is shown above. **b**, Structure of the human pre-A complex. The human complex was assembled on an exon-definition pre-mRNA. A schematic diagram of the pre-mRNA and its recognition by the pre-A complex is shown above.

Recently, small-molecule therapeutics targeting spliceosome have shown encouraging anti-cancer efficacy in clinical development ^60–62^. Most of these splicing modulators act on U2 snRNP ^10,21,63^. To gain structural insights, we incubated a representative splicing inhibitor E7107 with our purified sample of U2 snRNP and determined the structure. Overall structures of the inhibitor-free and E7107-bound U2 snRNP are almost identical, except that in the latter case E7107 fills the Hinged pocket ^38,39^ (Extended Data Fig. 14a, 14b). In our structure of the pre-A complex, SSA binds in the same general location but is covalently linked to the side chain of Cys26 from PHF5A (Extended Data Fig. 14c, 14d). Analysis of detailed interactions of the inhibitor with the pocket may facilitate development of potent splicing inhibitors. Notably, our observation confirms a published study in which SSA was reported to be covalently coupled to PHF5A of the SF3b complex ^31^.

PRP5 is anchored on SF3B1 through its N-terminal motifs; these interactions tether the RNA helicase domain of PRP5 to U2 snRNP. This general theme is reminiscent of the strategy employed by another essential RNA helicase Prp2.

Although Prp2 lacks such N-terminal motifs and is only loosely associated with the B^act^ complex, its coactivator Spp2 comprises four such motifs ^64^. Remarkably, two motifs of Spp2 tightly associate with the spliceosome, and the other two are anchored on Prp2. Using this strategy, Spp2 tethers Prp2 to the spliceosome. Future investigations on other ATPase/helicases will likely reveal similar but distinct strategies. Any such strategy must simultaneously allow stable association of the RNA helicase with the spliceosome and sufficient latitude of the helicase domain as necessitated by ATP binding and hydrolysis.

