## supplementary information for "Structural Insights into Branch Site Proofreading by Human Spliceosome"

### Methods

#### Preparation of human 17S U2 snRNP

Full-length PRP5 was subcloned into the pCAG expression vector with an N-terminal Flag tag. 1.5 mg of this plasmid was pre-incubated with 4 mg of PEIs in 50 mL of fresh medium at room temperature for 20 minutes. The resulting DNA-PEI mixture was added to one liter of HEK293 cell culture. At least four liters of cells were used for each preparation.

Cells were collected in the lysis buffer (20 mM HEPES-KOH, pH 7.9, 150 mM NaCl, 1.5 mM $MgCl_2$) supplemented with 0.05% NP-40 and disrupted by sonication. After removing cell debris and chromatin aggregates by centrifugation, the supernatant was applied to a FLAG affinity column. After extensive wash using the lysis buffer, proteins were eluted using the FLAG peptide. To remove excessive PRP5 and other contaminants, we further purified the sample using glycerol gradient centrifugation under crosslinking condition as described [1,2]. After centrifugation at 30,000 rpm for 20 hours at 4 °C in a SW32 rotor (Beckman Coulter), the sample was manually collected into 20 fractions from top to bottom. Total RNAs extracted from a 300-μl aliquot for each fraction were analyzed on an 8% denaturing polyacrylamide gel in the presence of 8 M urea. Based on results of the RNA gel, the fractions containing U2 snRNA only were pooled, concentrated using a 50-KDa cut-off Centricon (Millipore), and dialyzed against the lysis buffer using a 20-KDa Mini-lyzer (Pierce) for at least 5 hours. The final sample, which contains U2 snRNA and all known protein components of human U2 snRNP, was concentrated to 0.5 mg/ml for cryo-EM analysis.

### Design and preparation of the MINX-exon pre-mRNA

The pre-mRNA MINX-exon is derived from MINX [3] with the following modifications: the 5'-exon is removed and the 5'SS (GUAAGA) is placed just downstream of the 3'-exon. The resulting pre-mRNA construct (referred to as MINX-exon) comprises a 63-nt intron, a 39-nt exon, and a 5'SS. Three consecutive MS2-binding aptamers were introduced to the 3'-end of 5'SS and were used for MBP-MS affinity purification [4]. DNA templates for *in vitro* transcription were generated from PCR, and RNA substrates were prepared using T7 runoff transcription.

### *In vitro* splicing reaction

Nuclear extract was prepared from HeLa S3 cells essentially as described [5], except that the last centrifugation step after dialysis was omitted to avoid loss of splicing activity. *In vitro* splicing reactions were performed as described [6] using 20 nM pre-mRNA with 40% nuclear extract in the presence of 2 mM ATP, 20 mM creatine phosphate, 3 mM $MgCl_2$ and 60 mM KCl. When using spliceostatin A (SSA) to inhibit splicing reaction, 20 nM of the compound was pre-incubated with the nuclear extract on ice for 30 minutes before adding pre-mRNA to initiate the reaction.

### Preparation of the pre-A complex

For isolation of the pre-A complex, spliceosomes were assembled on the MINX-exon pre-mRNA substrate that had been incubated with MBP-MS2 protein in 200 mL *in vitro* splicing reaction supplemented with 20 nM SSA. After incubation at 30 °C for 60 minutes, large protein aggregates were removed through centrifugation and the assembled spliceosomes were purified using the amylose resin in a low-salt buffer (20 mM HEPES-KOH, pH 7.9, 75 mM NaCl, 4% glycerol and 1.5 mM $MgCl_2$). Eluent from the amylose resin was immediately applied to glycerol gradient centrifugation in the presence of 0-0.1% EM-grade glutaraldehyde as described [7], using a SW32 rotor (Beckmen) at 25,300 rpm. After 12 hours, 20 fractions of about 2 mL each from top to bottom were collected and the crosslinking reaction was quenched by addition of 50 mM Tri-HCl pH 7.5. The fractions containing the pre-A complex, as judged by RNA gel analysis, were pooled and concentrated using 100 kDa MWCO Amicon filters. The sample was dialyzed against the buffer (20 mM HEPES-KOH, pH 7.9, 75 mM NaCl, 1.5 mM $MgCl_2$) for 6 hours. The sample was analyzed on SDS-PAGE gels and further concentrated to 0.5 mg/mL for cryo-EM sample preparation and analysis.

**EM sample preparation and data acquisition**

To prepare cryo-EM specimen, the purified protein complexes were concentrated to 1.0 and 0.5 mg/ml for human 17S U2 snRNP and the pre-A complex, respectively. An aliquot of 3-μl sample was applied to a glow-discharged holey carbon grid (Quantifoil Au 300 mesh, R1.2/1.3), blotted for 3 seconds and rapidly plunged into liquid ethane using Vitrobot Mark IV (Thermo Fisher Scientific) operating at 8°C and 100% humidity.

The sample grids were imaged on a 300-kV Titan Krios electron microscope (Thermo Fisher Scientific) using a normal magnification of 81,000x. Movies were recorded using a Gatan K3 detector (Thermo Fisher Scientific) equipped with a GIF Quantum energy filter (slit width 20 eV) at the counted mode, with a pixel size of 0.5435 Å. Each stack of 32 frames was exposed for 2.56 seconds, with a dose rate of ~23 counts/second/physical-pixel (~19.5 e-/second/$Å^2$) for each frame. AutoEMation II [8] was used for the fully automated data collection with an efficiency of ~3000 stacks for each period of 24 hours. All 32 frames in each stack were aligned and summed using the whole-image motion correction program MotionCor2 [9] and binned to a pixel size of 1.087 Å. The defocus value for each image varied from 1.5 to 2.0 μm and was determined by Gctf [10].

**Cryo-EM data processing**

For 17S U2 snRNP, a total of 4,273 micrographs were collected, of which 3,606 micrographs were manually selected for further processing. A total of 1,972,935 particles were auto-picked using Gautomatch (https://www2.mrc-lmb.cam.ac.uk/research/locally-developed-software/zhang-software/#gauto) and subjected to 3D classification. The initial 3D volume of 17S U2 snRNP was generated from preliminary data analysis using cryoSPARC [11]. To avoid losing good particles, we simultaneously performed two parallel runs of single-reference 3D classification (Round 1). After Round 1, good particles were selected and merged, and the duplicated particles were removed. The remaining 847,852 particles were subjected to auto-refinement using the 2x binned particles (pixel size: 2.174 Å), resulting in a reconstruction at an average resolution of 4.4 Å (Round 2). Using the re-centered and re-extracted unbinned particles (pixel size: 1.087 Å), a reconstruction at an average resolution of 3.3 Å was generated. The remaining particles were classified (Round 3) with a soft mask on the core region of 17S U2 snRNP. The class containing 485,418 particles (57.2% of the input) yielded a reconstruction at an average resolution of 2.5 Å. These particles were further local classified and refined using different soft masks, generating four additional reconstructions for the PRP5-α1, PRP5-α2, TST-SF1 and BSL region at resolutions of 2.9 Å, 2.9 Å, 3.8 Å and 2.8 Å, respectively. In the 2.5-Å map of 17S U2 snRNP, the local resolution reaches 2.4 Å or even higher in the core region. The angular distributions of the particles used for final reconstruction of 17S U2 snRNP are reasonable, and refinement of the atomic coordinates does not suffer from severe over-fitting. The resulting EM density maps display clear features for amino acid side chains in the core region.

For the pre-A complex, a total of 22,451 micrographs were collected, of which 18,697 "good" micrographs were selected. A total of 5,156,919 particles were auto-picked using Gautomatch and were subjected to two parallel runs of single-reference 3D classification, followed by two parallel runs of local multi-reference 3D classification (Round 1). After Round 1, the remaining 2,197,019 particles were applied to two parallel runs of multi-reference 3D classification again (Round 2), but with 2x binned particles (pixel size: 2.174 Å). This step generated 1,466,041 good particles. Due to the relatively mobile nature of the interface between U1 and U2 snRNPs, we applied local masks on these two region for further 3D classification. Using the re-centered and re-extracted unbinned particles (pixel size: 1.087 Å), we generated reconstructions for U2 snRNP and U1 snRNP regions at average resolutions of 3.2 Å and 15.2 Å, respectively. These particles were further local classified and refined using different soft masks, generating two additional reconstructions for the U2 core region and DNAJC8 region both at resolutions of 3.0 Å, respectively.

Reported resolution limits were calculated on the basis of the FSC 0.143 criterion with a high-resolution noise substitution method [12]. Prior to visualization, all EM maps were corrected for modulation transfer function (MTF) of the detector, and then sharpened by applying a negative B-factor that was estimated using automated procedure [13]. Local resolution variations were estimated using ResMap [14] (Table S1).

**Model building and refinement**

We combined *de novo* model building and rigid docking of known structures to generate the atomic model (Tables S2 & S3). For human 17S U2 snRNP, the coordinates from a published study (PDB code: 6Y5Q) [15] were fitted into our 2.5-Å EM map and manually adjusted for each component. Higher resolution (2.4-3.0 Å) maps from focused refinement of the U2 snRNP core were used to guide model building in the core. The N-terminal portion of PRP5 (residues 152-243), which tethers PRP5 to SF3B1, was entirely *de novo* modeled on the basis of our EM map. This portion includes the helix α1, the acidic loop, and the helix α2. Notably, the N-terminal helix of PRP5 (α1), which is misplaced in the published study [15], has been unambiguously identified in our model. The RRM and Linker domain of TAT-SF1 were also *de novo* modeled on the basis of the EM map. In addition, a fragment of SF3B2 (residues 703-712) was found to bind SF3B3. The EM density for the small molecule E7107 shows fine details for its placement.

For the pre-A complex, the coordinates of human 17S U2 snRNP (this paper) and U1 snRNP (PDB code: 3CW1) [16] were docked into the EM maps for the U2 and U1 regions, respectively. For the U2 region, DNAJC8 was *de novo* built based on the EM map. The coordinates of the KH-QUA2 domain (PDB code: 1K1G) [17] and the coiled-coil domain of SF1 (PDB code: 4FXX) [18] were docked into the EM density maps and manually adjusted. The duplex between U2 snRNA and BS was built using the BSL from 17S U2 snRNP as a template.

The final models of human 17S U2 snRNP and the pre-A complex were refined on the basis of the EM maps using REFMAC in reciprocal space [19], with secondary structure restraints generated by ProSMART [20]. Overfitting of the model was monitored by refining the model in one of the two independent maps from the gold-standard refinement approach, and testing the refined model against the other [21]. The structures were validated through examination of the Molprobity scores and statistics of the Ramachandran plots (Table S1). Molprobity scores were calculated as described [22].

### *In vitro* binding assay between PRP5 and SF3B1

Missense mutations in SF3B1 and PRP5 were individually generated using sitedirected mutagenesis. The expression cassettes were individually subcloned into the pCAG vector with an N-terminal Flag and Twin-Strep tag. The binding assays were performed in two ways: either using purified wild-type (WT) Strep-PRP5 as the bait to pulldown various Flag-SF3B1 mutants, or conversely using purified WT Flag-SF3B1 to pulldown various Strep-PRP5 mutants. In the case of using WT Strep-PRP5 as the bait, 200 mL of cell culture were transfected to express WT or mutant Flag-SF3B1. Collected cells were lysed in 10 mL lysis buffer. 100 μg of purified Strep-PRP5 was added to the cell lysate. The mixture was incubated for 12 hours before loading into the Strep-Tactin resin. After extensive washing, proteins were eluted and analyzed by Western blots using an anti-Flag monoclonal antibody. The relative pulldown efficiencies as measured by the ratio of pulldown over input were averaged for three independent experiments and normalized to the corresponding WT protein. Conversely, in the case of using SF3B1 as the bait, WT or mutant Strep-PRP5 was expressed, incubated with Flag-SF3B1, purified through Flag-affinity resin, and analyzed using an anti-Strep monoclonal antibody.

**PRP5 RNA interference and RNA sequencing**

For knockdown of PRP5, HEK293T cells were maintained in Dulbecco's Modified Eagle Medium (DMEM) (Thermo Fisher Scientific) supplemented with 10% fetal bovine serum (FBS) (CellMax), 100 U/mL penicillin, and 100 μg/mL streptomycin (Hyclone) at 37 °C in a 5% $CO_2$ incubator. Chemically synthesized siRNAs targeting PRP5 and negative control siRNAs were purchased from GenePharma. Cells were transfected with siRNA at final concentration of 30 nM using Lipofectamine 2000 reagent (Thermo Fisher Scientific). 48 hours after transfection, total RNA and proteins were extracted using the TRIZOL reagent (Thermo Fisher Scientific) and RIPA lysis buffer (Cell Signaling Technology), respectively. Quantitative real-time PCR (qRT-PCR) and Western blots were performed to detect the knockdown efficiency of PRP5. All experiments were performed in triplicate. The sequences for siRNAs and primers are list in Table S4.

For RNA-sequencing, cDNA was synthesized with HiScript II 1st Strand cDNA Synthesis Kit (Vazyme) according to manufacturer's instructions, with quality assessments conducted on an Agilent 2100 RNA nano 6000 assay kit (Agilent Technologies). Sequencing libraries were generated using VAHTS Universal V6 RNA-seq Library Prep Kit for Illumina (NR604-01/02) following manufacturer's recommendations and index codes were added to attribute sequences to each sample. The clustering of the index-coded samples was performed on a cBot cluster generation system using HiSeq PE Cluster Kit v4-cBot-HS (Illumina) according to manufacturer's instructions. After cluster generation, the libraries were sequenced on an Illumina platform (Novaseq 6000 S4) and 150 bp paired-end reads were generated.

**RNA-Seq analysis and validation**

Alternative splicing analysis was carried out using JuncBASE [23]. Briefly, sequencing reads were mapped to human hg19 genome index with STAR (v2.7.7a) (10.1093/bioinformatics/bts635). The STAR index files were generated using hg19 genome sequence and Gencode (10.1101/gr.135350.111) annotation file (Ensembl 88 GRCh37). The generated bam files by STAR were sorted using Samtools (v1.3.1) (10.1093/bioinformatics/btp352). Then JuncBASE was used to identify and quantify alternative splicing events and perform differential splicing analysis between WT and PRP5 knockdown cells. The statistically significant differential alternative splicing events were defined as those with $|\Delta PSI|>10$ and P-value<0.05 (Table S5).

Quantitative Taqman assays were used to detect the normal and aberrant spliced transcripts as described [24]. Relative splicing activity was measured by calculating the ratio of aberrant to normal spliced forms for each target gene. All primers used are list in Extended Data Table 4.

**Data availability:** The atomic coordinates for human 17S U2 snRNP and the pre-A complex have been deposited in the Protein Data Bank (PDB) under the accession codes 7EVO and 7VPX, respectively. The EM maps of 17S U2 snRNP core region, PRP5-α1, PRP5-α2, TAT-SF1 and BSL have been deposited in the EMDB with accession codes EMD-31334, EMD-31335, EMD-31336, EMD-31337 and EMD-31338, respectively. The EM map of the pre-A complex core region, DNAJC8 region and U1 snRNP region have been deposited in EMDB with accession codes EMD-32074, EMD-32075, and EMD-32076, respectively. The accession number for the RNA sequencing data reported in this paper is GEO: GSE185713.

**Acknowledgements**

We thank the Cryo-EM Facility, the Computing Center, the Protein Characterization and Crystallography Facility and the Mass Spectrometry & Metabolomics Core Facility of Westlake University for technical support. We thank Nicholas A. Larsen from H3 Biomedicine for providing SSA.

**Funding**

This work was supported by funds from the National Natural Science Foundation of China (31930059 to Y.S.), the China Postdoctoral Science Foundation (2020M671806 to X. Zhang, 2021M692888 to X. Zhan), the National Postdoctoral Program for Innovative Talents of China (BX20200305 to X. Zhang, BX2021268 to X. Zhan), and Start-up funds from Westlake University (to Y.S.).

**Author Contributions**

Y.S. conceived the project. X. Zhang, T.B., and F.Y. designed and performed the experiments. X. Zhang prepared cryo-EM samples and collected the EM data. X. Zhan and X. Zhang processed the EM data, calculated the EM maps, and built the atomic models. F.Y., P.L. and Q.Z. performed RNA-seq data analysis. Y.S. and X. Zhang wrote the manuscript with input from all authors.

**Competing interests**

The authors declare no competing financial interests.

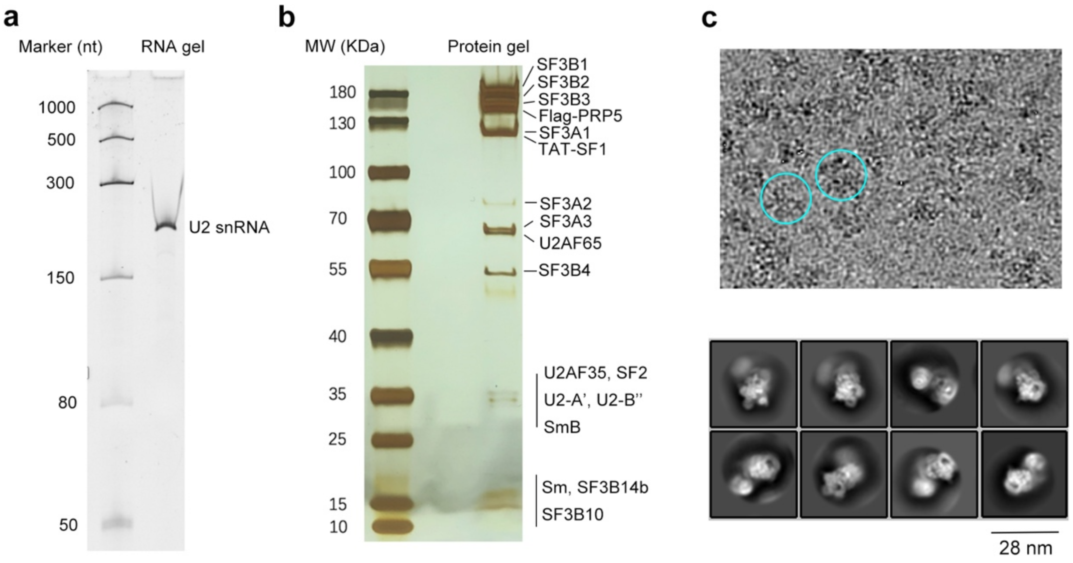


**Extended Data Fig. 1 Purification and preliminary characterization of human 17S U2 snRNP.** **a**, Only U2 snRNA is detectable in the purified human 17S U2 snRNP as shown in this denaturing PAGE gel. **b**, Analysis of the purified human 17S U2 snRNP on a silver-stained SDS-PAGE gel. The protein components were identified by mass spectrometry. **c**, A representative cryo-EM micrograph (upper panel) and representative 2D class averages (lower panel) of the human 17S U2 snRNP sample.

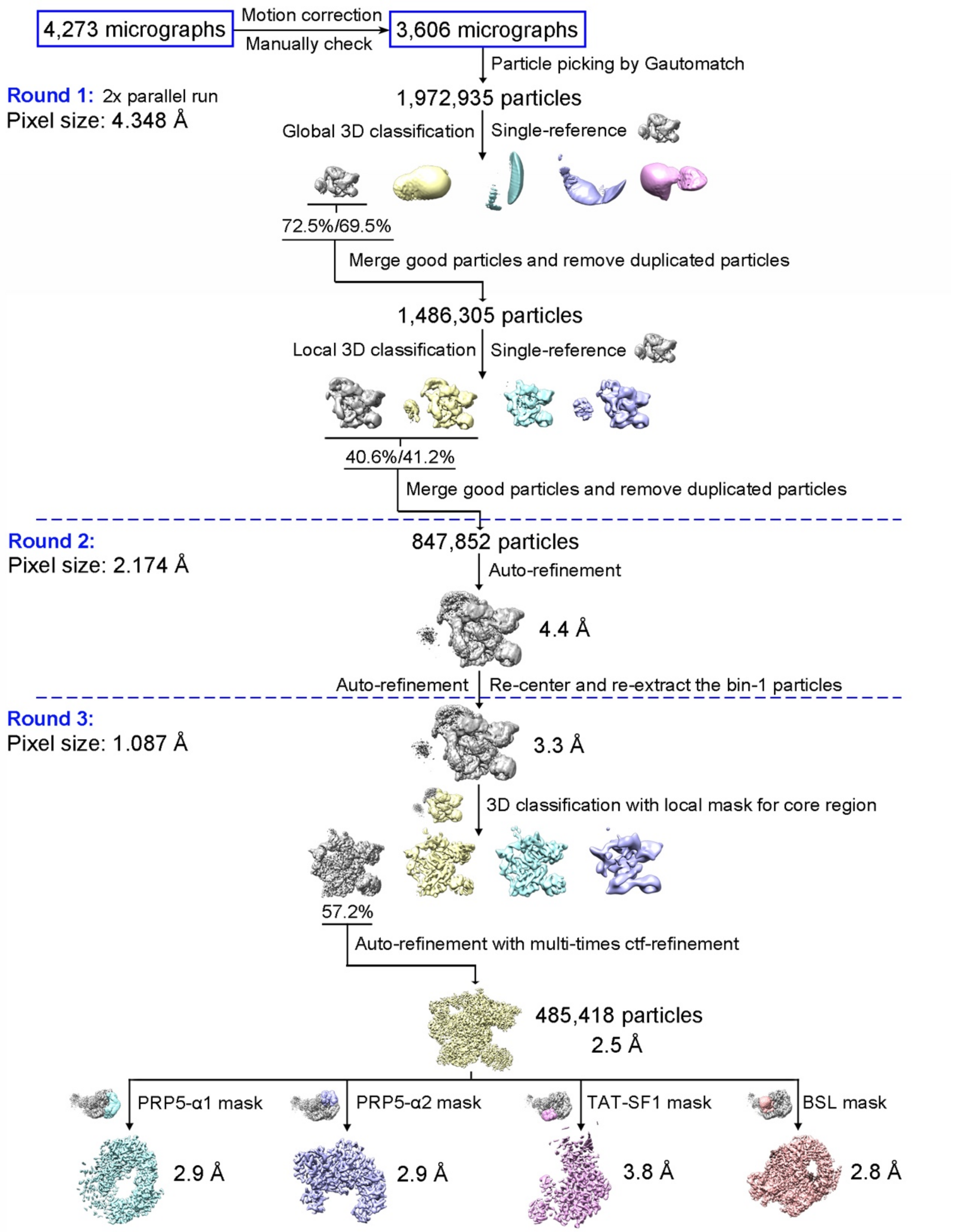


**Extended Data Fig. 2 A flow chart diagram of cryo-EM data processing for human 17S U2 snRNP.** All processing steps were carried out in RELION 3.0 [25]. Please refer to Methods for details.

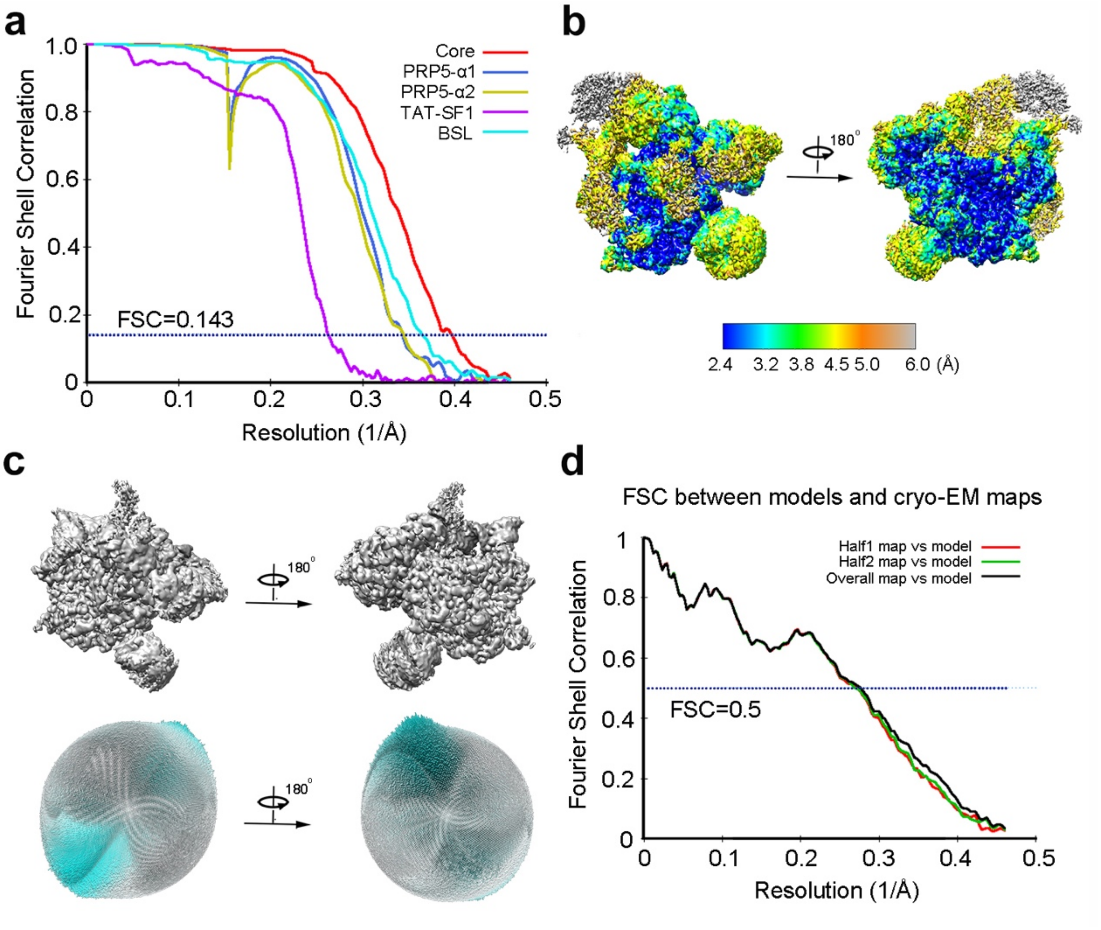


**Extended Data Fig. 3 Cryo-EM reconstruction of human 17S U2 snRNP at an average resolution of 2.5 Å.** **a**, The final reconstructions for the masked core (red), PRP5-α1 (blue), PRP5-α2 (yellow), TAT-SF1(magenta) and BSL (cyan) display average resolutions of 2.5 Å, 2.9 Å, 2.9 Å, 3.8 Å and 2.8 Å, respectively, on the basis of the FSC value of 0.143. **b**, Two overall views of the EM density map. The local resolutions are color-coded for different regions of U2 snRNP. **c**, Angular distribution of the particles used for the reconstruction. Each cylinder represents one view and the height of the cylinder is proportional to the number of particles for that view. **d**, The Fourier-shell correlation (FSC) curves for cross-validation between the model and the cryo-EM map of human 17S U2 snRNP. Shown here are the FSC curves between the final refined atomic model and the reconstruction from all particles (black), between the model refined in the reconstruction from only half of the particles and the reconstruction from that same half (red), and between that same model and the reconstruction from the other half of the particles (green).

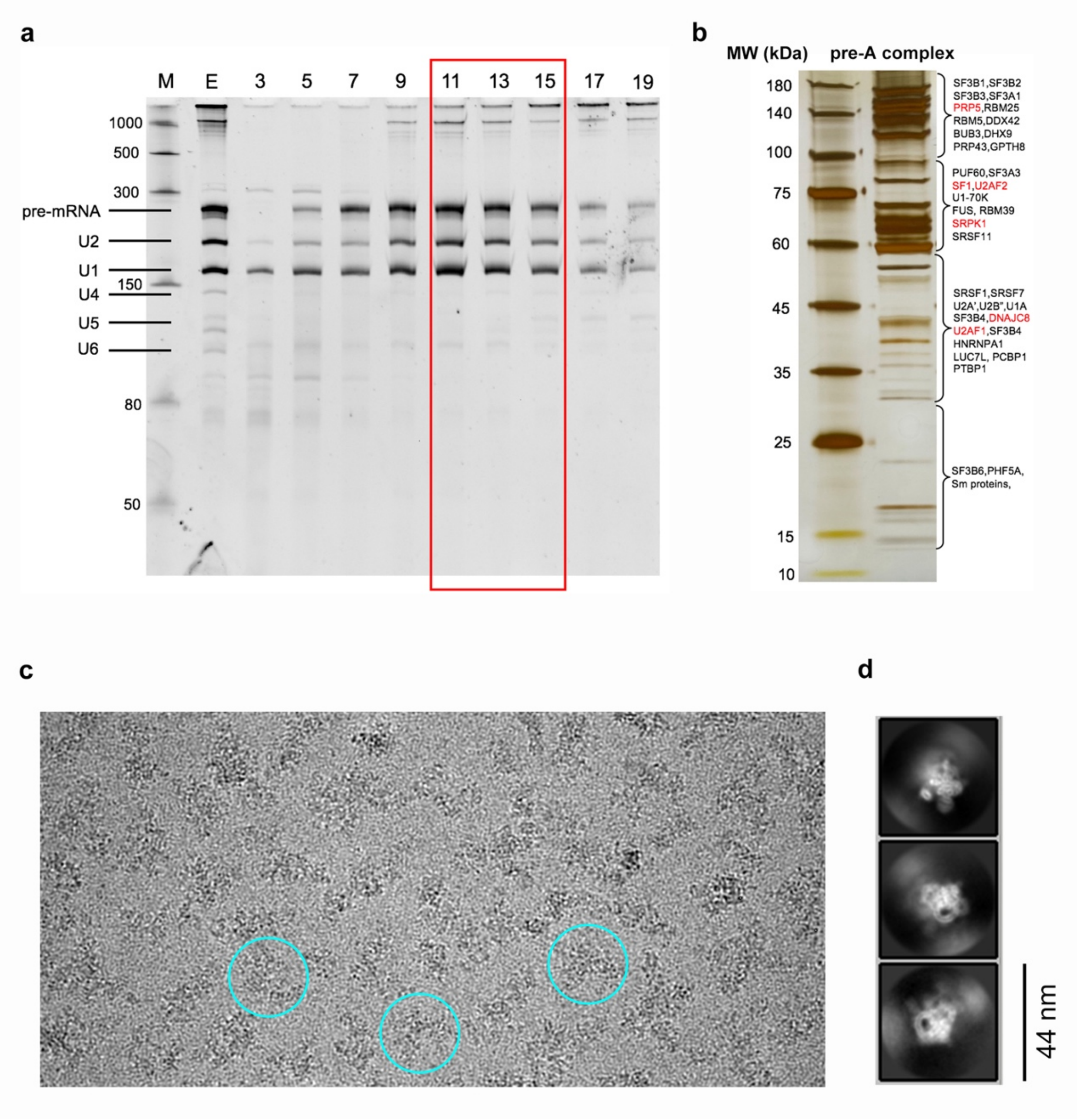


**Extended Data Fig. 4 Purification and preliminary characterization of the human pre-A complex.** **a**, Analysis of RNA components on 8% urea-PAGE gel after glycerol gradient centrifugation. The fractions that contain pre-mRNA and five snRNAs are indicated by a red rectangular box. Fractions 11-15, thought to contain the pre-A complex, were pooled for cryo-EM analysis. **b**, Analysis of the purified pre-A complex on a silver-stained SDS-PAGE gel. The protein components were identified by mass spectrometry. **c**, A representative cryo-EM micrograph of the pre-A complex sample. **d**, Representative 2D class averages. Scale bar: 44 nm.

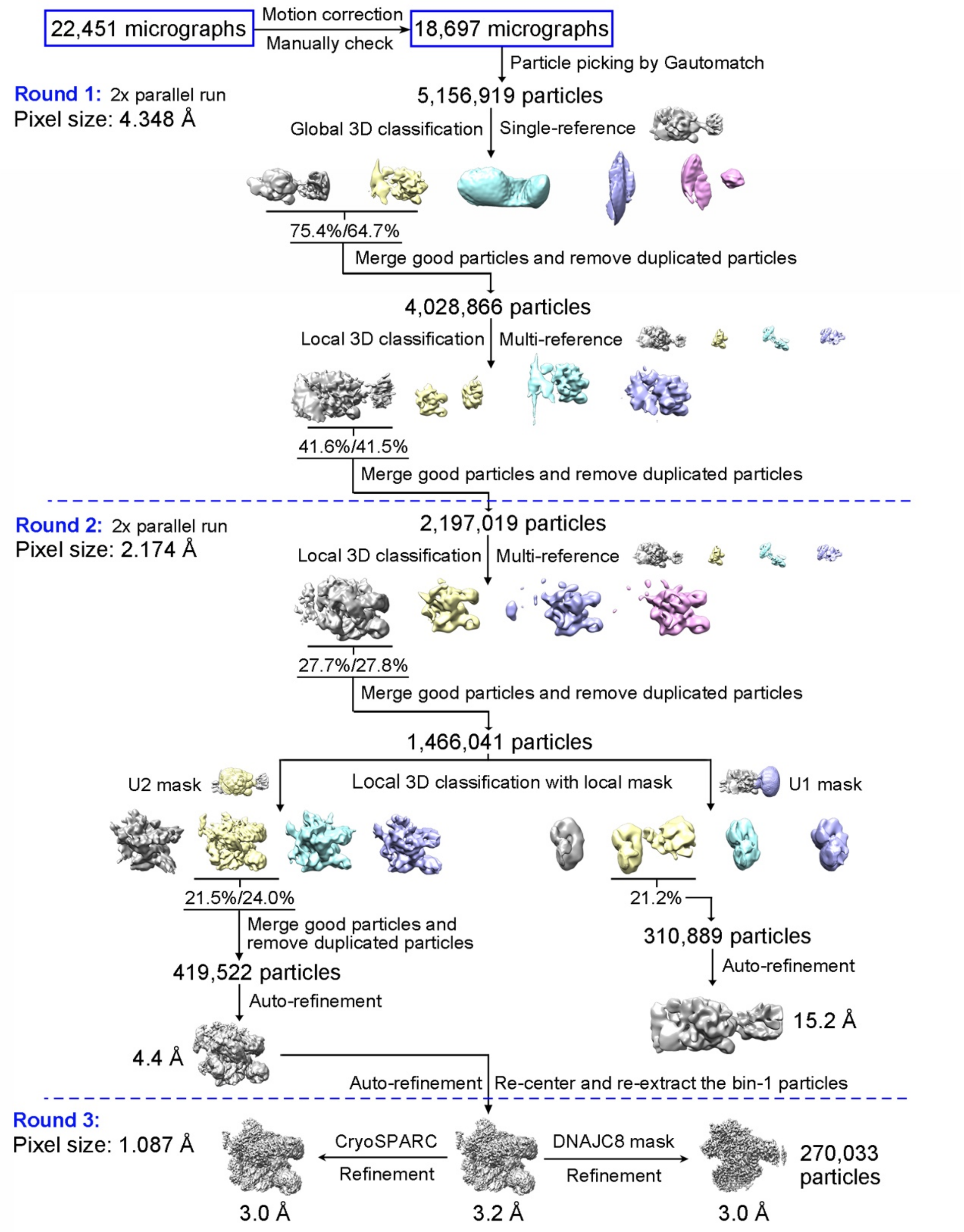


**Extended Data Fig. 5 A flow chart diagram of cryo-EM data processing for the human pre-A complex.** All processing steps were carried out in RELION 3.0 [25]. Please refer to Methods for details.

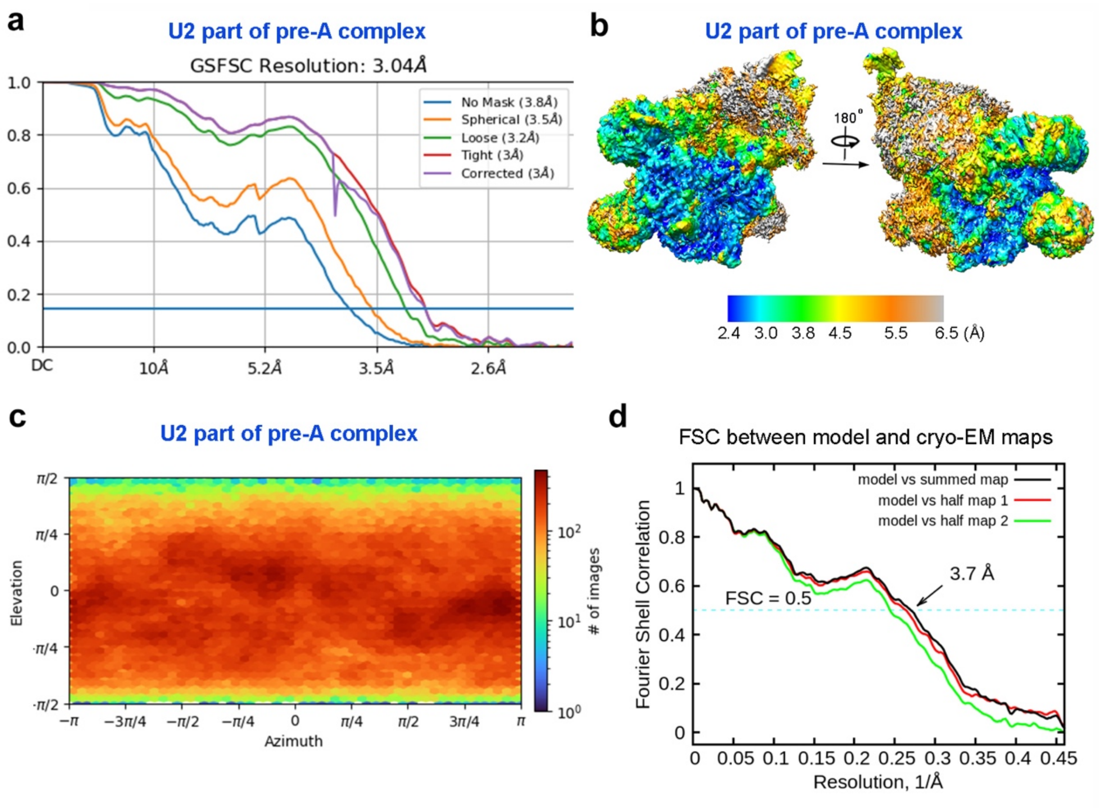


**Extended Data Fig. 6 Cryo-EM reconstruction of the U2 snRNP region of human pre-A complex at an average resolution of 3.0 Å.** **a**, The final reconstruction for the masked U2 region displays an average resolution of 3.0 Å on the basis of the FSC value of 0.143. **b**, Two overall views of the EM density map. The local resolutions are color-coded for different regions of U2 snRNP in the pre-A complex. **c**, Angular distribution of the particles used for the reconstruction of U2 snRNP region of human pre-A complex. **d**, The Fourier-shell correlation (FSC) curves for cross-validation between the model and the cryo-EM map for the human pre-A complex. Shown here are the FSC curves between the final refined atomic model and the reconstruction from all particles (black), between the model refined in the reconstruction from only half of the particles and the reconstruction from that same half (red), and between that same model and the reconstruction from the other half of the particles (green).

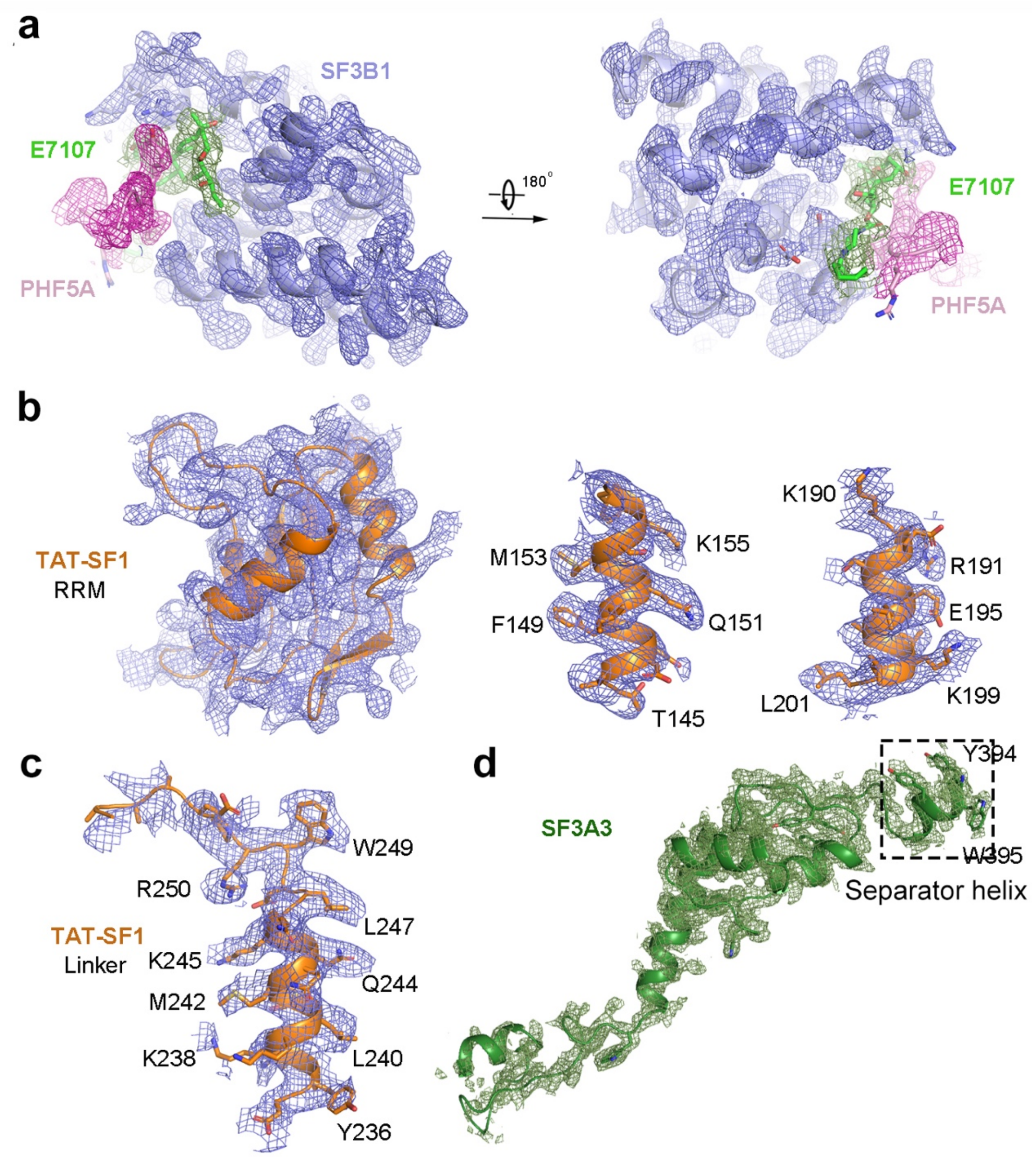


**Extended Data Fig. 7 The EM density map of human 17S U2 snRNP.** **a**, The EM density map for the small molecule E7107 and its surrounding structural elements. Two related views are shown. **b**, The EM density map for the RRM domain of the splicing factor TAT-SF1 (left panel). The EM density maps for two representative α-helices are shown (middle and right panels). **c**, The EM density map for the Linker domain of TAT-SF1. **d**, The EM density map for SF3A2. The separator helix is indicated by a black dashed box.

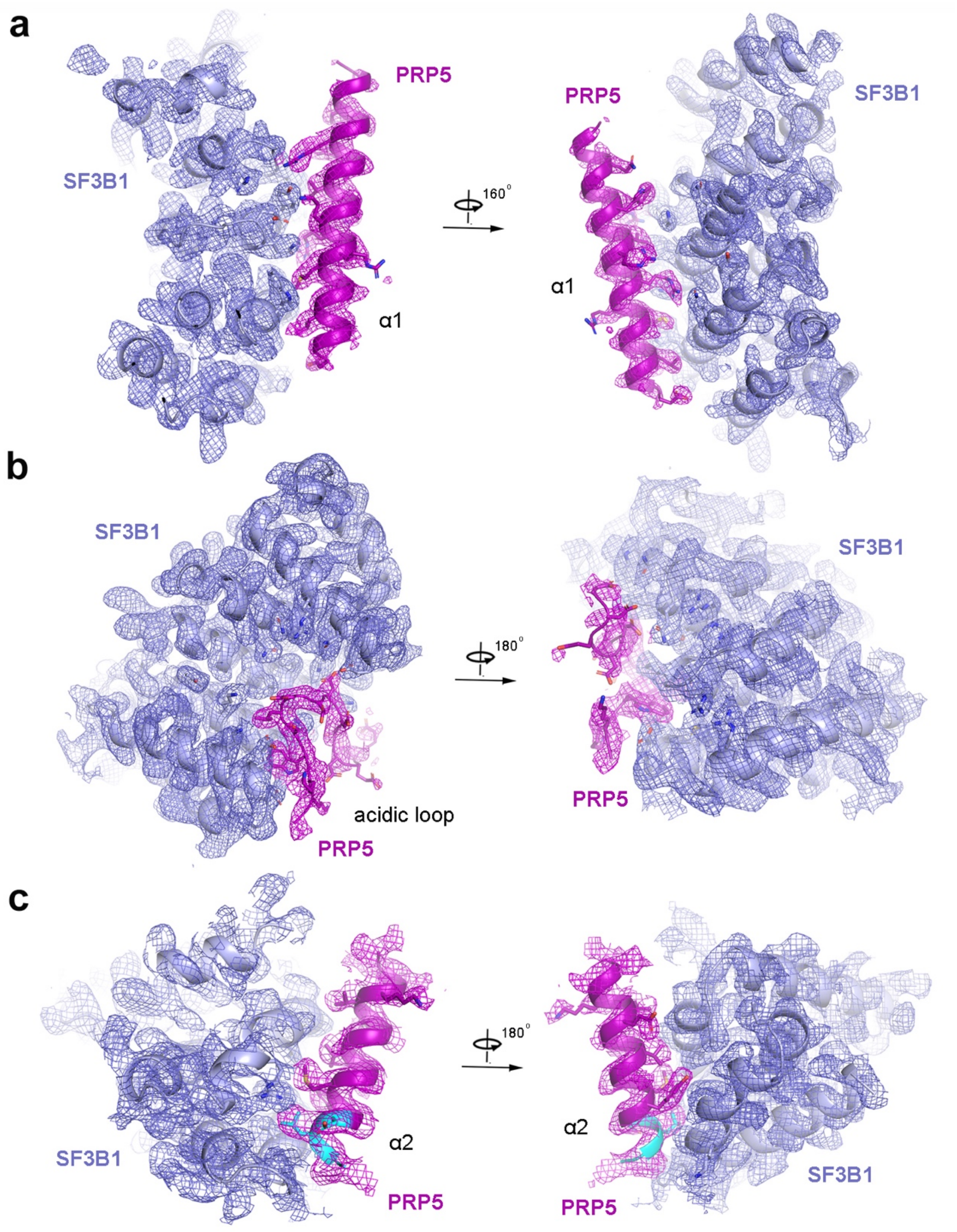


**Extended Data Fig. 8 The EM density map for the sequences of PRP5.** **a**, The EM density map for the α1 helix of PRP5 and its interacting elements from SF3B1. Two related views are shown. **b**, The EM density map for the acidic loop of PRP5 and its interacting elements from SF3B1. Two related views are shown. **c**, The EM density map for the α2 helix of PRP5 and its interacting elements from SF3B1. Two related views are shown.

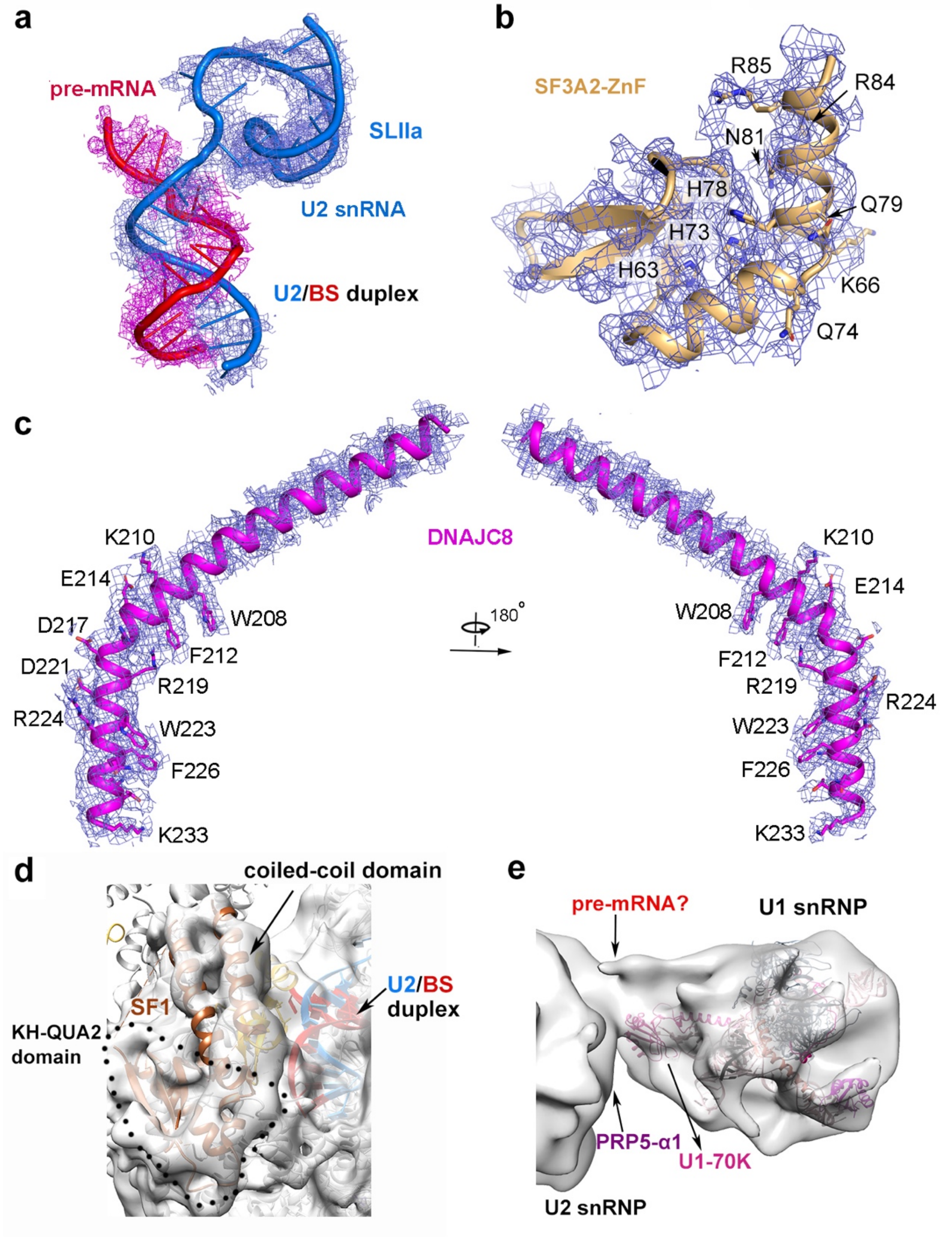


**Extended Data Fig. 9 The EM density map for different regions of the human pre-A complex.** **a**, The EM density map for the initial U2/BS duplex in the pre-A complex. **b**, The EM density map for the zinc finger (ZnF) of SF3A2. **c**, The EM density map for the structurally resolved region of DNAJC8. Two related views are shown. **d**, Docking of SF1 into the EM density map. **e**, Docking of U1 snRNP into the EM density map.

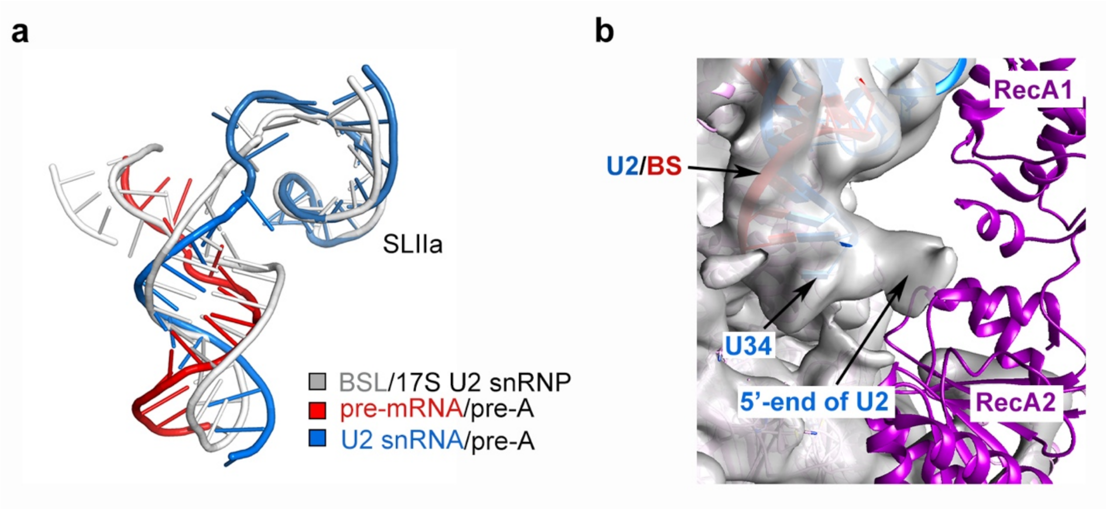


**Extended Data Fig. 10 Formation of an initial U2/BS duplex in pre-A complex.** **a**, Superimposition of the initial U2/BS duplex and the three-way junction of U2 snRNA from human 17S U2 snRNP (grey). **b**, The low-pass filtered density map for the 5'-end sequences of U2 snRNA can be traced to the interface between RecA1 and RecA2 of PRP5.

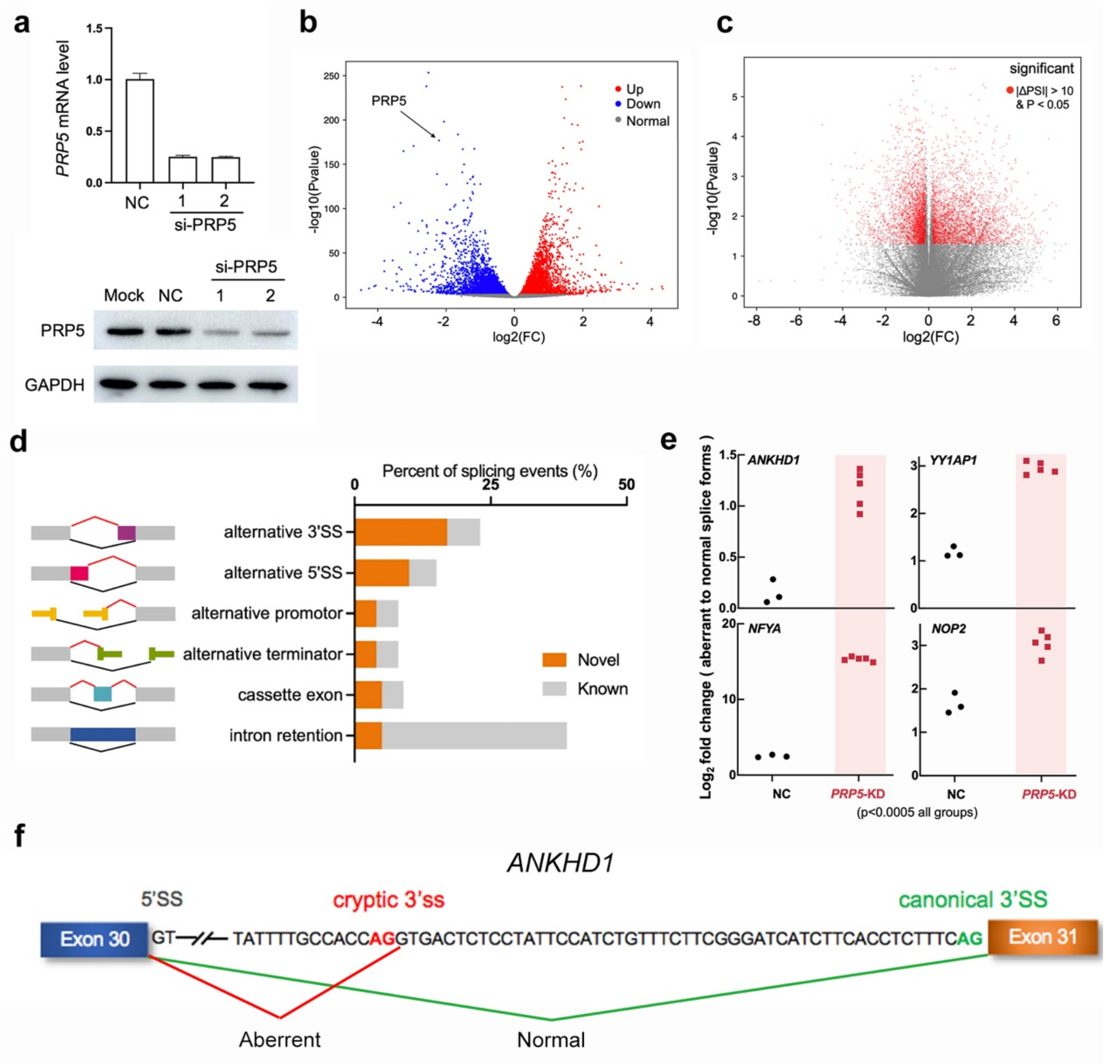


**Extended Data Fig. 11 PRP5 knockdown in HEK293 cells induces altered splicing site selection.** **a**, Analysis of *PRP5* mRNA level (top panel) and protein expression level (bottom panel) in control (NC) and siRNA-transfected cells. **b**, The volcano plot of $\log_2$ fold change versus $\log_{10}$(Pvalue) of gene expression level in siRNA-transfected cells and WT cells. Red and blue dots indicate up-regulated and down-regulated genes, respectively (Pvalue < 0.001). **c**, The volcano plot of $\log_2$ fold change of ΔPSI versus $\log_{10}$(Pvalue) of all splicing changes. Red dots indicate significantly altered splicing events (|ΔPSI| > 10 and Pvalue < 0.05). **d**, Distribution of novel (orange) and known (grey) aberrant splicing events in PRP5 knockdown cells. Intron retention and alternative 3'SS are the most frequently occurring events. **e**, Validation of representative aberrantly spliced genes in control (n=3) and PRP5 knockdown (n=5) cells through quantitative RT-PCR assays. **f**, A schematic diagram of normal and aberrant splicing of ANKHD1, a representative gene that is sensitive to PRP5 knockdown. Positions of the cryptic (red) and canonical (green) 3'SS are indicated.

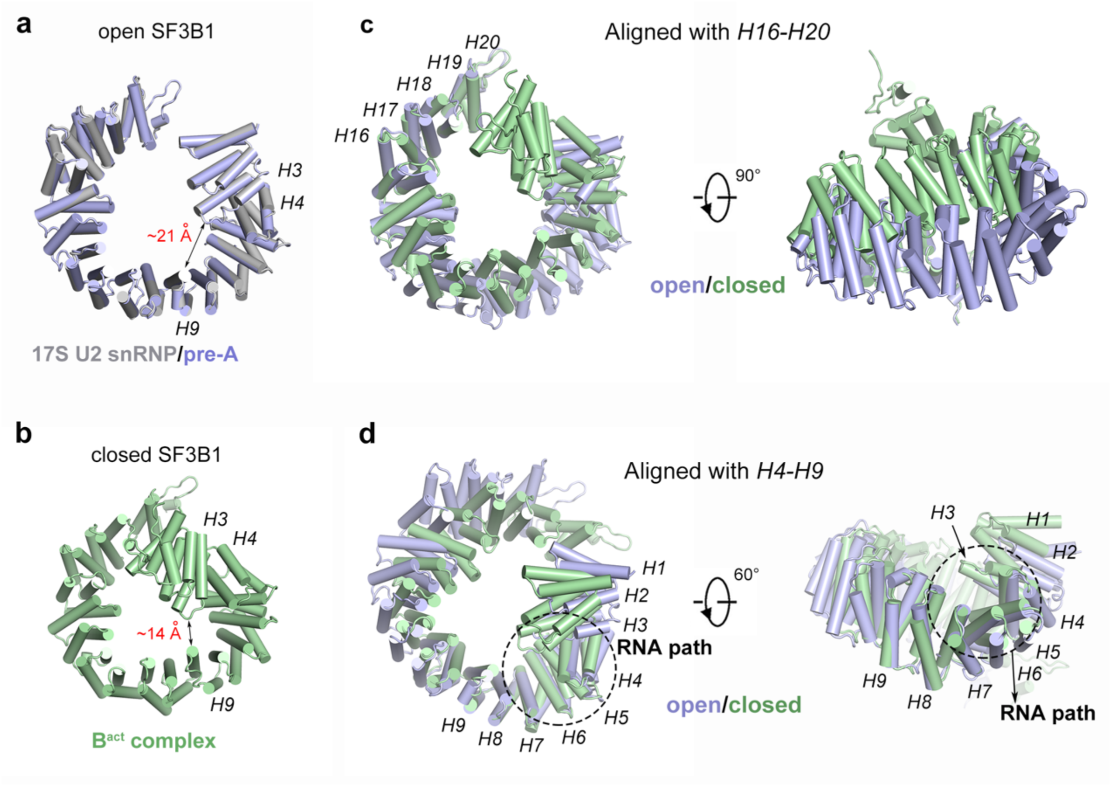


**Extended Data Fig. 12 The open and closed conformations of SF3B1.** **a**, The open conformation of SF3B1. SF3B1 exists in an open conformation in both 17S U2 snRNP and the pre-A complex. Shown here is an overlay of the SF3B1 structures from these two complexes. **b**, The closed conformation of SF3B1. Pre-mRNA engagement by U2 snRNP results in a closed conformation of SF3B1. Shown here is the structure of SF3B1 from the human $B^{act}$ complex [26]. **c**, Structural alignment using HEAT repeats 16 through 20 of SF3B1 reveals major conformational differences between the open and closed states of SF3B1. Two perpendicular views are shown. **d**, Structural alignment using HEAT repeats 4 through 9 of SF3B1 reveals major conformational differences between the open and closed states of SF3B1.

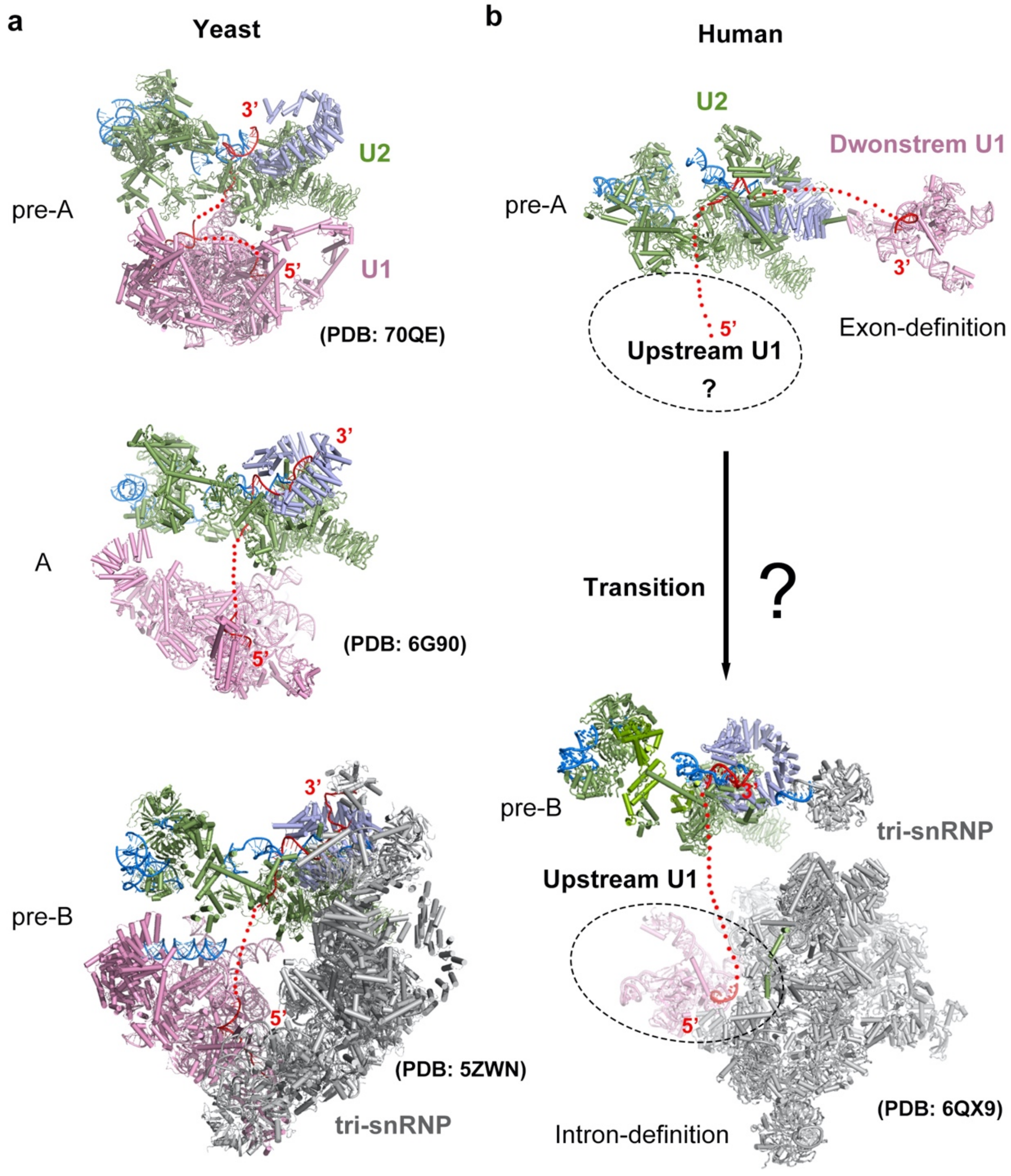


**Extended Data Fig. 13 The relative orientations of U1 and U2 snRNPs in yeast and human spliceosomes.** **a**, Molecular organizations of yeast U1 (pink) and U2 (green and light blue) snRNPs in the pre-A, A and pre-B complexes. Overall, the U1 snRNP moves away from U2 snRNP to allow the recruitment of tri-snRNP. **b**, In human, the cross-exon pre-A complex should be converted to cross-intron complex for splicing reaction to proceed; this step may involve interaction of U2 snRNP with another U1 snRNP that bound an upstream 5’SS.

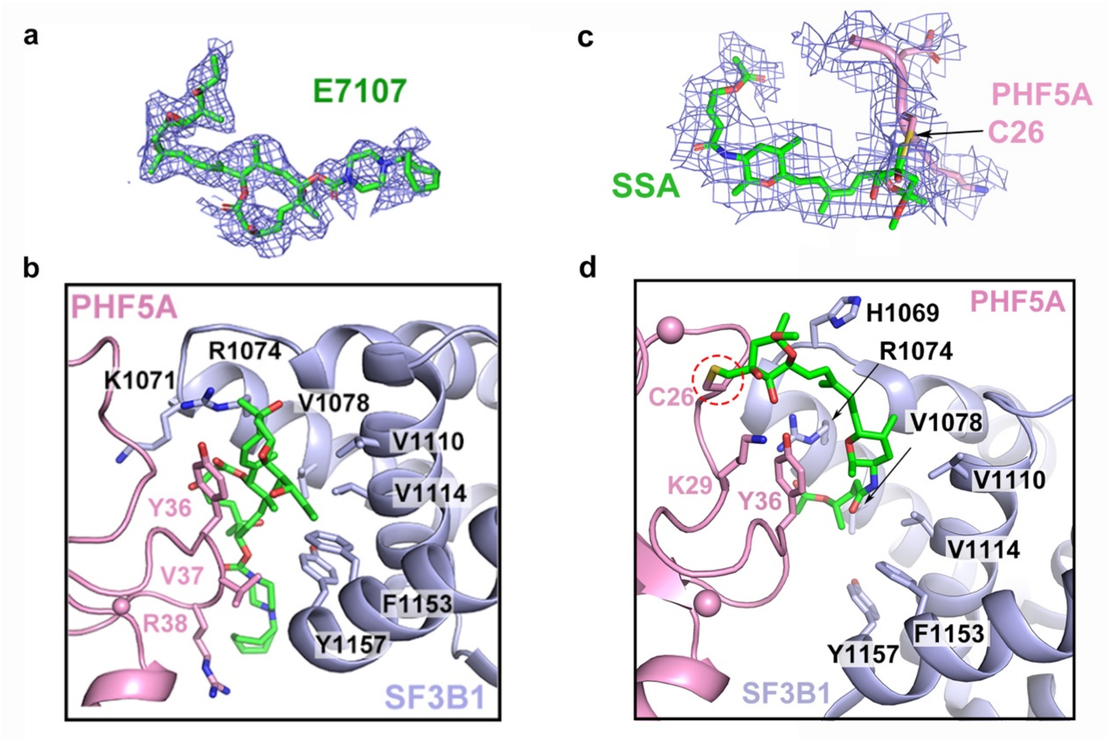


**Extended Data Fig. 14 Recognition of the splicing inhibitors E7107 and SSA by the Hinged pocket.** **a**, A close-up view on the EM density of E7107 in human 17S U2 snRNP. **b**, A close-up view on the interface between E7107 and the Hinged pocket. The Hinged pocket is exactly where the BPA binds in the assembled spliceosomes. **c**, A close-up view on the EM density of SSA in the human pre-A complex. SSA is covalently linked to Cys26 of PHF5A. **d**, A close-up view on the interface between SSA and the Hinged pocket.

**Extended Data Table 1. Statistics of EM analysis and model validation.**

| | 17S U2 snRNP | pre-A complex |
|---|---|---|
| **Data collection** | | |
| EM equipment | FEI Titan Krios | |
| Voltage (kV) | 300 | |
| Detector | K3 | |
| Pixel size (Å) | 1.087 | |
| Electron dose (e-/Å$^2$) | 50 | |
| Defocus range (μm) | 1.5~2.0 | |
| **Reconstruction** | | |
| Software | RELION 3.0 | |
| EMDB code | EMD-31334 | EMD-32074 |
| Number of particles | 485,418 | 419,522 |
| Symmetry | C1 | |
| Final masked resolution (Å) | 2.5 | 3.0 |
| Map sharpening B-factor (Å$^2$) | -80.4 | -100.1 |
| **Model building** | | |
| Software | Coot 0.8.9 | |
| Refinement | Phenix/Refmac | |
| PDB code | PDB: 7EVO | PDB: 7VPX |
| Protein residues | 4871 | 5612 |
| RNA nucleotides | 148 | 319 |
| **Validation** | | |
| R.m.s deviations | | |
| Bonds length (Å) | 0.01 | 0.01 |
| Bonds Angle (°) | 1.75 | 1.76 |
| Ramachandran plot statistics (%) | | |
| Preferred | 94.30 | 91.79 |
| Allowed | 5.22 | 7.37 |
| Outlier | 0.48 | 0.84 |
| CaBLAM outliers (%) | 3.0 | 2.6 |
| MolProbity score | 2.79 | 2.66 |

### Extended Data Table 2. Summary of model building for human 17S U2 snRNP.

| | Molecule | Length | Domain/Region | PDB code | Modeling | Resolution (Å) | Chain ID |
|---|---|---|---|---|---|---|---|
| **17S U2 snRNP** | U2 snRNA | 188 nt | 12:73/81:184 | 6Y5Q | RD | 3.5~8.0 | H |
| | SF3B1 | 1304 | 490:1304 | | RD | 2.5~3.0 | 1 |
| | SF3B2 | 895 | 458:598<br>703:712 | 6Y5Q<br>/ | RD<br>DM | 2.5~3.0 | 2 |
| | SF3B3 | 1217 | 1:1217 | 6Y5Q | RD | 2.5~3.0 | 3 |
| | SF3B4 | 424 | 11:181 | | RD | 6.0~10.0 | 4 |
| | SF3B5 | 86 | 16:81 | | RD | 2.5~3.0 | 5 |
| | PHF5A | 110 | 6:91 | | RD | 2.5~3.0 | 6 |
| | SF3A1 | 793 | 160:282 | | RD | 6.0~10.0 | A |
| | SF3A2 | 464 | 104:209 | | RD | 6.0~10.0 | B |
| | SF3A3 | 501 | 1:501 | | RD | 6.0~10.0 | C |
| | U2-A’ | 255 | LRR domain | | RD | 6.0~10.0 | F |
| | U2-B” | 225 | RRM domain | | RD | 6.0~10.0 | G |
| | SmB,D1,D2<br>D3,E,F,G | - | Sm fold | | RD | 6.0~10.0 | a-g |
| | TAT-SF1 | 755 | 157:256<br>UHM (264:347) | /<br>6Y5Q | DM<br>RD | 3.0~4.0<br>5.0~8.0 | D |
| | PRP5 | 1031 | 152:243<br>helicase domain | /<br>6Y5Q | DM<br>RD | 2.5~3.5<br>6.0~8.0 | E |
| | E7107 | - | - | - | DM | 2.5~3.0 | 1 |

Under the column labeled “Modeling”, DM stands for *de novo* modeling; RD stands for rigid docking and manual adjustment.

**Extended Data Table 3. Summary of model building for the human pre-A complex.**

| | Molecule | Length | Domain/Region | PDB code | Modeling | Resolution (Å) | Chain ID |
|---|---|---|---|---|---|---|---|
| | U2 snRNA | 188 nt | 35:65/70:184 | 6Y5Q | RD/DM | 3.5~8.0 | H |
| | Pre-mRNA | - | branch sequence | / | DM | 3.5~4.5 | I |
| | SF3B1 | 1304 | 490:1304 | 6Y5Q | RD | 3.0~3.5 | 1 |
| | SF3B2 | 895 | 458:598<br>703:712 | 6Y5Q<br>/ | RD<br>DM | 3.0~3.5 | 2 |
| | SF3B3 | 1217 | 1:1217 | | RD | 3.0~3.5 | 3 |
| | SF3B4 | 424 | 11:181 | | RD | 6.0~10.0 | 4 |
| | SF3B5 | 86 | 16:81 | | RD | 3.0~3.5 | 5 |
| | PHF5A | 110 | 6:91 | | RD | 3.0~3.5 | 6 |
| | SF3A1 | 793 | 160:282 | | RD | 6.0~10.0 | A |
| **U2 snRNP** | SF3A2 | 464 | 41:87/104:209 | 6Y5Q | RD | 4.0~10.0 | B |
| | SF3A3 | 501 | 1:501 | | RD | 6.0~10.0 | C |
| | U2-A’ | 255 | LRR domain | | RD | 6.0~10.0 | F |
| | U2-B” | 225 | RRM domain | | RD | 6.0~10.0 | G |
| | SmB,D1,D2<br>D3,E,F,G | - | Sm fold | | RD | 6.0~10.0 | a-g |
| | SF1 | 639 | 46:119<br>134:229 | 4FXX<br>1K1G | RD | 5.0~8.0 | D |
| | PRP5 | 1031 | 152:243<br>helicase domain | /<br>6Y5Q | DM<br>RD | 3.5~4.5<br>6.0~8.0 | E |
| | DNAJC8 | 253 | 180:233 | / | DM | 3.5~5.0 | J |
| | SSA | - | - | - | DM | 3.0~3.5 | 6 |
| | U1 snRNA | 164 nt | 1:164 | | RD | 10.0~15.0 | L |
| | Pre-mRNA | - | 5’SS | | RD | 10.0~15.0 | K |
| | U1-A | 282 | 6:103 | | RD | 10.0~15.0 | N |
| **U1 snRNP** | U1-C | 159 | 2:51 | 3CW1 | RD | 10.0~15.0 | M |
| | U1-70K | 437 | 2:202 | | RD | 10.0~15.0 | O |
| | SmB,D1,D2<br>D3,E,F,G | - | Sm fold | | RD | 10.0~15.0 | h-n |

Under the column labeled “Modeling”, DM stands for *de novo* modeling; RD stands for rigid docking and manual adjustment.

**Extended Data Table 4. Sequences of siRNAs and primers.**

| siRNAs and primers | Sequences |
|---|---|
| **siRNAs** | |
| Negative control siRNA | 5'-UUCUCCGAACGUGUCACGUTT-3' |
| PRP5 siRNA | 5'-GCAGAAGCTGAAAAGGAG-3' |
| **primers for Taqman assay** | |
| GAPDH (Forward) | 5'-AAATCCCATCACCATCTTCC-3' |
| GAPDH (Reverse) | 5'-AGGCTGTTGTCATACTTCTC-3' |
| PRP5 (Forward) | 5'-CGATGATGATGACGAAGATG-3' |
| PRP5 (Reverse) | 5'-CTCCTCTGAAGAATACTCCAT-3' |
| ANKHD1 (Forward) | 5'-TCTCCCCATCCTTGGACAAG-3' |
| ANKHD1 (Normal Reverse) | 5'-ACTGACTCCTTTTGGTTGGC-3' |
| ANKHD1 (Aberrant Reverse) | 5'-TGAAAGAGGTGAAGATGATC-3' |
| ANKHD1 (Normal probe) | 5'-AACTCATCCACTTCTGC-3' |
| ANKHD1 (Aberrant probe) | 5'-ACTCATGTGACTCTCCTA-3' |
| NFYA (Forward) | 5'-TCAACCAGTTAATGCAGATG-3' |
| NFYA (Normal Reverse) | 5'-AATCTGTGCTCCTGCCAAAC-3' |
| NFYA (Aberrant Reverse) | 5'-TGCTGGGATAGTGATCATGC-3' |
| NFYA (Normal probe) | 5'-TCCAGCAAGGCATGATC-3' |
| NFYA (Aberrant probe) | 5'-CAGCAAGTTACAGTCCCTG-3' |
| NOP2 (Forward) | 5'-CTGAGGCCAAACCATTGC-3' |
| NOP2 (Reverse) | 5'-ATTAAATAGGGACTGGGGTC-3' |
| NOP2 (Normal probe) | 5'-AGCTACCAAAAGGAGCTGT-3' |
| NOP2 (Aberrant probe) | 5'-AAAGGGATCTCTGCAGGAG-3' |
| YY1AP1 (Forward) | 5'-CAGGATAAGATCCTCTTCAC-3' |
| YY1AP1 (Normal Reverse) | 5'-TTCTCACTGTCAGTTGGCG-3' |
| YY1AP1 (Aberrant Reverse) | 5'-TTAAGAAACTCAGTCCCTTC-3' |
| YY1AP1 (Normal probe) | 5'-CTGAGGACAACAAGTACCT-3' |
| YY1AP1 (Aberrant probe) | 5'-AGGACAATTTGTTAGCTT-3' |
